# Maternal diet alters offspring’s early life host-microbiota communication through goblet cells, resulting in long-lasting diseases susceptibility

**DOI:** 10.1101/2024.07.05.602179

**Authors:** Clara Delaroque, Erica Bonazzi, Marine Huillet, Sandrine Ellero-Simatos, Fuhua Hao, Andrew Patterson, Benoit Chassaing

**Author notes:** Corresponding Authors: Benoit Chassaing, Ph.D. Institut Pasteur, Université Paris Cité, INSERM U1306, *Microbiome-Host Interaction* group Paris, France. **Abbreviations:** CMC : Carboxymethylcellulose; P80 : polysorbate 80.

## Abstract

A crucial early-life developmental phase regulates microbiome settling while establishing critical and long-lasting immune and metabolic processes. During this period, the influence of select components of maternal diet on offspring microbiota and health remains largely unknown. To investigate the potential transgenerational impact of maternal exposure to microbiota-disrupting factors, dams were subjected prior breeding to dietary emulsifiers, known to directly perturb the microbiota. Such maternal exposure induced early-life microbiota alterations in offspring which associated with long-lasting susceptibility to diet-induced obesity and intestinal inflammation. These detrimental effects were entirely prevented by early-life microbiota normalization through cross-fostering procedures. Mechanistically, maternal emulsifier exposure induces strong offspring’s impairment in goblet cells-mediated host-microbiota communication which is central in driving the observed long-lasting deleterious effects. To conclude, this study underscores the central role played by maternal intake of microbiota-disrupting agents on the next generation’s microbiota, with long lasting consequences for intestinal and metabolic health.

## Introduction

At birth, individuals encounter intestinal colonization by a highly diverse range of microbial species, leading to settlement of the intestinal microbiota. Such microbial exposure during a specific time window framed by birth and weaning drives immune system activation, facilitating the development and maturation of immune cells and organs ^1–8^, thus establishing a symbiotic relationship between the host and its microbiota. Disruption of this primo colonization during early life results in immunological and metabolic unreversible defects that persist into adulthood ^1,4^, suspected to lead to predisposition to various diseases including inflammatory diseases ^1,9,10^, asthma ^11,12^, cancer ^1^ and metabolic deregulations ^2,13–15^. One proposed explanation for the importance of this critical time window involves the early-life specific circulation of microbiota-derived antigens through colonic goblet cells (goblet cells-associated antigens passage, GAPs) ^16–18^, which was reported to be central in driving immune tolerance towards commensals ^16,18,19^.

Recognizing the pivotal role played by early-life microbiota in adulthood host health, research efforts have focused on identifying environmental factors influencing the establishment of this microbial community. Environmental factors such as early-life antibiotic treatments ^2,20^, maternal influence through transmission of select bacterial strains ^21,22^, as well as indirect mechanisms including the involvement of milk components and genetic factors ^23–25^, have emerged as a contributors of early-life microbiota settlement. However, the exact human relevance of these findings, as well as the underlying long-lasting mechanisms at play, both remain unclear. Hence, we investigated here whether maternal exposure to broadly consumed microbiota stressors could influence offspring’s early-life microbiota in a way that will subsequently impact adulthood’s intestinal and metabolic health. For this purpose, we applied exposure to dietary emulsifier, additives commonly added in ultra-processed foods, as a model of microbiota perturbation. Recent literature reported that dietary emulsifiers are directly impacting the intestinal microbiota and interact with the epithelial lining in a way that favor chronic intestinal inflammation and metabolic deregulations, in both mice and humans ^26–29^. Moreover, recent epidemiological data from the *NutriNet Santé* cohort, including >175,000 participants, reported positive associations between intake of dietary emulsifiers and risk of cardiovascular diseases, various cancers as well as type 2 diabetes^30–32^.

Here, we observed that offspring from emulsifier-exposed dams exhibited early-life microbiota alterations, at both the compositional and functional levels, which associated with increased adulthood susceptibility to diet-induced obesity and colitis. Microbiota normalization through cross-fostering procedures revealed the causal relationship between early-life microbiota alterations and long-lasting disease susceptibility. Mechanistically, offspring from emulsifier-exposed dams displayed anticipated GAPs closure, while chemical prevention of such GAPs closure was sufficient to fully prevent long-lasting susceptibility to intestinal inflammation and metabolic deregulations. Altogether, these data suggest that maternal exposure to common dietary factors is sufficient to prevent proper early-life host-microbiota interactions, with a central role played by the intestinal microbiota and goblet cells homeostasis, in a way that drive transgenerational long-lasting susceptibility to inflammation and metabolic deregulations.

## Results

### Maternal emulsifier consumption induces early-life alterations in offspring’s microbiota composition

To investigate the trans-generational effect of dietary emulsifiers consumption on early life microbiota, C57Bl/6J dams were subjected to 1% of carboxymethylcellulose (CMC) or polysorbate-80 (P80) in drinking water for 10 weeks prior breeding (**Figure 1a**). Such treatment was sufficient to reproduce previously observed microbiota alteration in dams ^26,29^ (**Fig. S1**). After 10 weeks of treatment, breeding pairs (N=6 per experimental group) were formed with unexposed male, and pregnant females were individually housed from gestation until weaning. At birth, offspring were either kept with their biological mother (experimental groups Water; CMC; P80) or cross fostered to a water-treated dam with age-matching and size-matching litters (experimental groups Water→Water; CMC→Water; P80→Water), with 3-7 litters used per experimental group. Male offspring were weaned at 3 weeks under emulsifiers free conditions, as described **Figure 1a**. While dams were kept under emulsifier treatment during the pre-weaning phase, we confirmed that offspring were not directly exposed to dietary emulsifiers through milk consumption, with the observation that CMC and P80 were both undetectable by NMR in milk (data not shown) ^28^.

**Figure 1.**
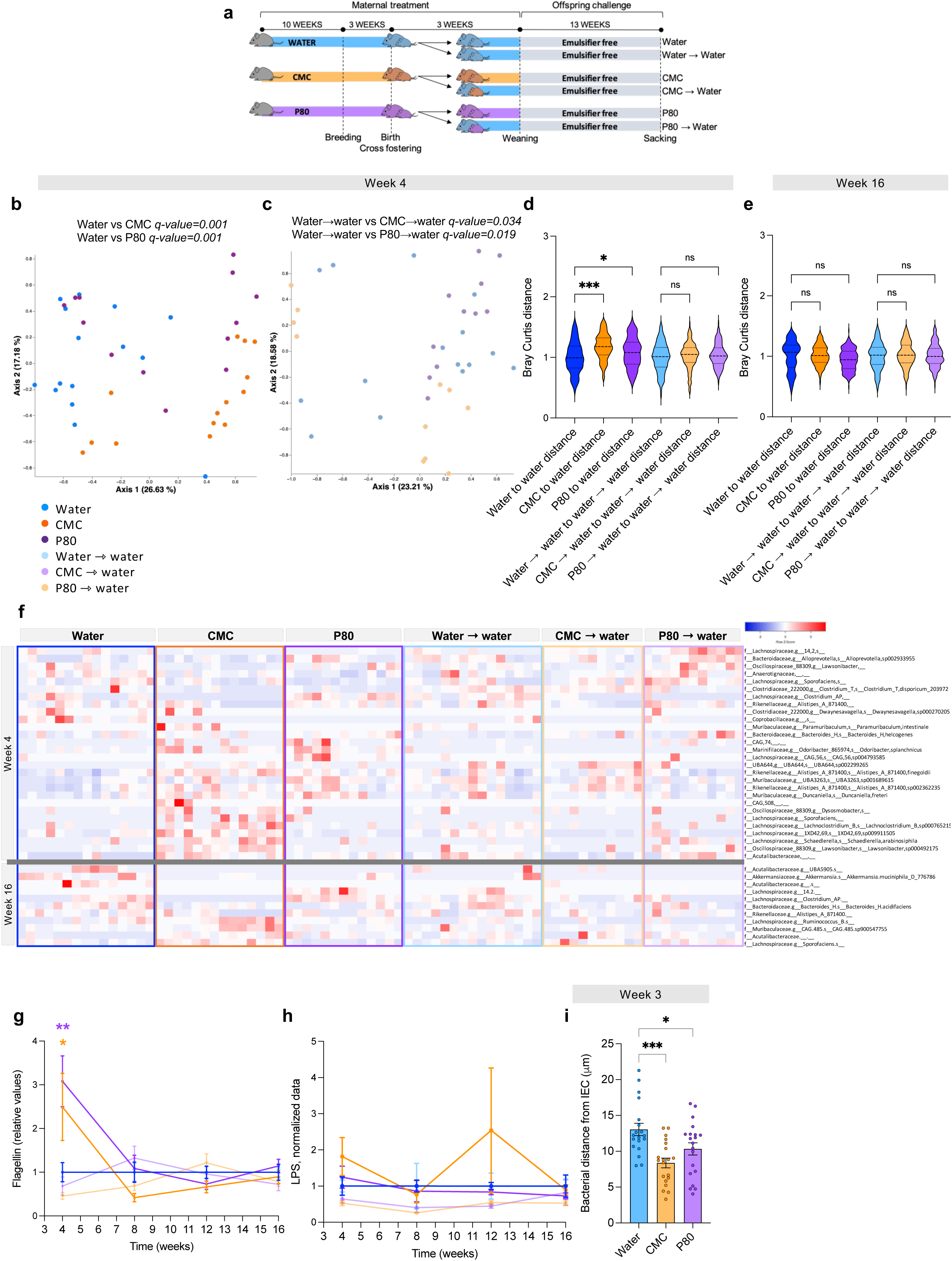
Maternal emulsifier consumption induces early-life functional and compositional microbiota alterations in offspring. Dams were subjected to 1% CMC or P80 in their drinking water for 10 weeks prior to breeding. At birth, offspring were either kept with their biological mother or cross-fostered to a water-treated dam. Following weaning of male offspring under emulsifier-free conditions, longitudinal compositional and functional analysis of the offspring’s microbiota were performed. (**a**) Schematic representation of the experimental design used. (**b-c**) Principal coordinates analysis of the Bray–Curtis distance at week 4, representing offspring from Water, CMC-, and P80-treated dams (**b**), or Water→Water, CMC→Water, and P80→Water cross-fostered offspring (**c**). (**d-e**) Bray-Curtis distance separating the various experimental groups from the water or Water→Water offspring at week 4 (**d**) and at week 16 (**e**). (**f**) Relative abundance of MaAsLin2-identified microbiota members with significantly altered abundance in CMC and P80 offspring compared to control, at week 4 and week 16. (**g-h**) Fecal levels of bioactive flagellin (**g**) and LPS (**h**), with data normalized to either Water or Water→Water offspring. (**i**) Distance of closest bacteria to intestinal epithelial cells (IECs) at week 3, over five high-powered fields per mouse. Data are presented as means ± s.e.m (N = 13-15 from ≥ 3 independent litters). Significance was assessed by ANOSIM (**b-c**), one-way ANOVA (**d, e, i**) or two-way ANOVA (**g, h**), and is indicated as follow: n.s. indicates non-significant * p≤ 0.05; **p ≤ 0.01; ***p ≤ 0.001.

Offspring’s fecal microbiota composition was next investigated by 16S rRNA gene sequencing. At 4 weeks of age, the overall microbiota composition was significantly divergent between offspring from water-, CMC- and P80-treated dams (**Fig. 1b, d**). Moreover, such alteration in composition was prevented when pups were cross-fostered by a water-treated dam (**Fig. 1c, d**). In adulthood, at 16 weeks of age, no significant alteration in microbiota composition was observed between CMC, P80 and water offspring (**Fig. 1e**), suggesting that the observed trans-generational alterations are restricted to the early-life period. Machine learning approach to predict dams’ treatment based on offspring microbiota composition revealed a loss of prediction accuracy throughout time (**Fig. S2 a-h**), further highlighting that microbiota alterations in offspring from emulsifier-treated dams are occurring in early-life before normalizing at adulthood.

We next performed Microbiome Multivariable Associations with Linear Models (MaAsLin2)-based analysis to identify microbiota members significantly altered in their abundance in CMC and P80 offspring compared to water offspring at week 4 or 16. Such an approach revealed 28 impacted microbiota members at week 4, compared to only 11 at week 16 (**Fig. 1f**) which, together with the absence of alteration in caecal-derived metabolites between groups in adulthood (**Fig. S2i, j**), further highlight that trans-generational alterations in microbiota composition are vanishing during adulthood.

Various mechanisms are at play in shaping early-life microbiota, including direct mom-to-infant microbiota members transmission. In addition, milk components are known to influence the developing microbiota in offspring ^23,25^, such as milk-derived IgA since mice pups start to produce their own IgA only after weaning, making maternal milk as the only source of IgA in early life ^24^. CMC offspring displayed a significant reduction in the proportion of IgA-bound bacteria, which was fully prevented by cross fostering (**Fig. S3a**). While we observed modest modification in select antigen-specific milk-derived IgA, such as anti-LPS IgA (**Fig. S3b-d**), this was not sufficient to significantly impact the overall bacterial community targeted by these antibodies (**Fig. S3e**). We next investigated if milk-derived bacteria are impacted by emulsifier consumption in a way that can drive the observed early life microbiota alterations in the descendants. 16S rRNA sequencing-based approach revealed similar composition in bacterial community in milk from CMC- or P80-treated compared to unexposed dams (**Fig. S3f, g**). We next assessed maternal milk metabolic profile, known to be a player in offspring’ microbiota establishment ^33,34^. NMR-based quantification identified 17 metabolites with a similar overall profile (**Fig. S3h, i**), and only modest emulsifier-induced alterations in alanine (**Fig. S3j**), creatine (**Fig. S3k**) and betaine (**Fig. S3l**) being observed. Altogether, these data indicate that maternal emulsifiers intake induces transient transgenerational microbiota alteration in composition, which may result from a combination of direct mom-to-pups microbiota transmission, together with modest alterations in milk-derived IgA and metabolites.

### Maternal emulsifier consumption induces early-life alterations in offspring’s intestine-microbiota interactions

We next investigated the potential trans-generational impact of emulsifier consumption on the offspring’s microbiota at the functional level. For this purpose, we first quantified microbiota-derived pro-inflammatory molecules lipopolysaccharide (LPS) and flagellin, proxi markers of a given microbiota pro-inflammatory potential ^35,36^ and previously reported to be significantly increased upon emulsifier exposure ^26,29,37^. Longitudinal fecal levels of bioactive flagellin, but not LPS, displayed significant increase at week 4 in CMC and P80 offspring compared to water offspring (**Fig. 1g, h**). Such observation indicates that offspring from emulsifier-treated dams display early-life restricted microbiota with increased pro-inflammatory potential, which can be fully prevented by cross fostering procedure (**Fig. 1g, h**).

With the previous observation that microbiota pro-inflammatory potential positively correlate with microbiota encroachment phenomenon ^26,38–40^, defined as the ability of select microbiota members to penetrate the normally sterile inner mucus layer, we next quantified microbiota-epithelium distance at weaning (**Fig. 1i**). Such distance was significantly reduced in CMC and P80 offspring compared to control offspring, which could be explain, at least in part, by the observed increased level of bacterial motility factor flagellin (**Fig. 1g**) and impaired IgA-mediated mucosa extrusion (**Fig. S3b-d**). These data further suggest early life transgenerational detrimental impact of maternal emulsifier consumption on the offspring’s intestinal microbiota, at both the compositional and functional levels.

### Maternal emulsifier consumption induces long-lasting susceptibility to diet-induced obesity

We next investigate whether such early-life trans-generational microbiota alteration induced by maternal emulsifiers intake could impact host metabolism. For this purpose, males weaned under emulsifiers-free conditions were subjected to a high-fat diet (HFD) regimen for 13 weeks, as presented **Fig. 2a**. Such an approach importantly revealed that CMC and P80 offspring gained significantly more weight upon HFD regimen compared to water controls (**Fig. 2b**). Moreover, early-life microbiota normalization through cross-fostering procedure was sufficient to prevent such increased body weight gain (**Fig. 2b**). Furthermore, offspring from P80-treated dams additionally presented increased peri-epididymal fat pad weight (**Fig. 2c**) as well as fasting glycemia (**Fig. 2d**), which were both prevented by cross fostering. Based on the major impact of HFD regimen on hepatic health, we next investigated liver homeostasis by histology and RT-q-PCR. While no differences were observed in liver weight between groups (**Fig. S4a**), major HFD-induced liver alterations were observed in CMC and P80 offspring compared to water offspring, as evidenced by significantly increased steatosis (**Fig. 2e** and **S4b**), early-stage fibrosis evidenced by Sirius red staining (**Fig. 2f**) and altered expression of steatosis-associated genes *Acta2 Pparg* and *Il1b* (**Fig. 2g**, **Fig. S4c, d**). In addition to liver damages, HFD regimen is also known to induce chronic low-grade intestinal inflammation^41^. Histological scoring of colonic sections revealed that CMC and P80 offspring displayed exacerbated HFD-induced low-grade intestinal inflammation (**Fig. 2h**), further evidenced by colon shortening and increased weight, both markers of colonic inflammation and immune cells infiltration (**Fig. S4e, f**). Importantly, all these HFD-induced perturbations observed in CMC and P80 offspring were fully abrogated in water cross-fostered offspring, thus revealing that transgenerational impact of maternal emulsifier consumption have potent and long-lasting deleterious impact on metabolic health, despite only transient detrimental effect on offspring’s intestinal microbiota.

**Figure 2.**
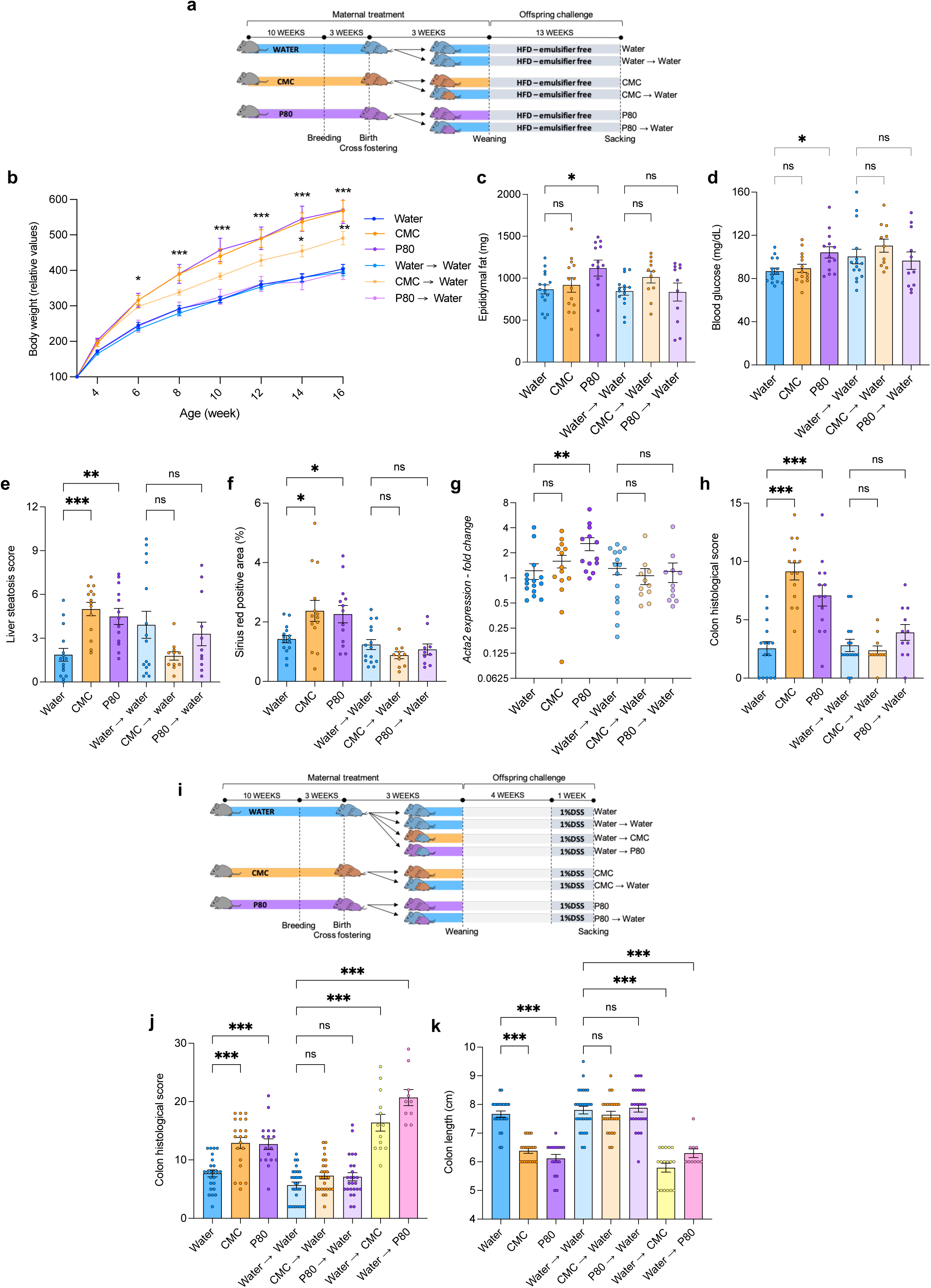
Maternal emulsifiers consumption induces long-lasting susceptibility to diet-induced obesity and colitis in offspring. Dams were subjected to 1% CMC or P80 in their drinking water for 10 weeks prior to breeding. At birth, offspring were either kept with their biological mother or cross-fostered to a water-treated dam. Following weaning of male offspring under emulsifier-free conditions, susceptibility to metabolic deregulations and intestinal inflammation was assessed by subjecting mice to a HFD regimen or DSS treatment, respectively. (**a**) Schematic representation of the experimental design used to investigate susceptibility to HFD-induced metabolic deregulations. (**b**) Body weight over time, with data being expressed as percentage compared to the body weight at weaning (week 3). (**c**) Epididymal fat deposition measured at sacking, (**d**) 15h fasting glycemia at week 14. After sacking, liver histological samples were stained with HE or Sirius red, allowing (**e**) steatosis histological scoring and (**f**) fibrosis quantification as the percentage of sirus red positive area. (**g**) Liver expression fold change of *Acta2.* (**h**) Histopathological scoring of HE stained colon samples following HFD challenge. (**i**) Schematic representation of the experimental design used to investigate susceptibility to DSS-induced colitis. (**h**) Histopathological scoring of HE stained colon samples following DSS challenge and (**k**) colon length. Data are the means ± s.e.m (N = 10-28 from ≥ 3 independent litters). Significance was assessed by two-way ANOVA (**b**) or one-way ANOVA (**c-h, j, k**) and is indicated as follow: n.s., non-significant; * ≤ 0,05; **p ≤ 0,01; ***p ≤ 0,001.

### Maternal emulsifier consumption induces long-lasting susceptibility to colitis

In light of the central role played by the early-life intestinal microbiota on long lasting susceptibility to intestinal inflammation ^1,2,13^, we next investigated colitis susceptibility in offspring from CMC- and P80-treated dams. For this purpose, males weaned under emulsifier-free conditions were subjected, 4 weeks after weaning (at 7 weeks of age), to 1% dextran sodium sulfate (DSS) diluted in the drinking water for 7 days, as presented **Fig. 2i**. Such an approach revealed that offspring from CMC- and P80-treated dams had an increased DSS-induced intestinal inflammation, as evidenced by an increased colonic histopathological score and colon shortening (**Fig. 2j, k**). CMC and P80 offspring also harbored an increased colon weight (**Fig. S5a**), suggesting immune cells infiltration, further evidenced by an increased infiltration of CD68-positive immune cells (**Fig. S5b**). Such increased colitis susceptibility in in CMC and P80 offspring was associated with alteration in mucus layer homeostasis, as revealed by decreased *Muc2* gene expression (**Fig. S5c**), together with alterations in select colonic and ileal lamina propria immune cell clusters (**Fig. S6**). These alterations were compound specific, aligning with previous observations on CMC and P80 ^26,27^. Moreover, early life microbiota normalization through cross-fostering procedure in CMC→water and P80→water groups (**Fig. 2i**) fully prevented the increased colitis susceptibility normally observed in CMC and P80 offspring, suggesting that early-life microbiota alterations are drivers of colitis susceptibility (**Fig. 2j, k and Fig. S5a, b**). Next, and thanks to the use of a relatively low dose of DSS (1%), we investigated metabolic health in these groups by quantifying peri-epididymal fat deposition. Well aligned with the above observations under HFD regimen, this revealed an increased adiposity in CMC and P80 offspring compared to water control offspring, which was fully prevented though cross-fostering by untreated dams (**Fig. S5d**).

Finally, to assess whether emulsifier consumption-induced transgenerational early life microbiota alterations are sufficient to drive colitis susceptibility, another cross-fostering approach was applied in which pups from water-treated dams were either kept with their own mother or cross foster to a water (water→water group), CMC-(water→CMC group) or P80- (water→P80 group) treated dams, as presented in **Fig. 2i**. Such cross-fostering approach was sufficient to drive early life microbiota alteration, as presented **Fig. S5e**, together with a stark increased in colitis susceptibility, compared to water→water control group, in adulthood (**Fig. 2j, k and Fig. S5a**). Altogether, these data highlight that early life microbiota alterations induced by maternal emulsifier intake are both necessary and sufficient to drive long-lasting increased susceptibility to colitis.

### Early-life microbiota alterations resulting from maternal exposure to dietary emulsifiers induce impaired goblet cells-associated passage at weaning

The observation of transgenerational susceptibility to metabolic deregulations and colitis despite microbiota normalization in adulthood prompted us to focus on the early-life time window, previously described as having long-lasting impact on disease susceptibility ^1,2,42^. Hence, we next investigated the effect of maternal emulsifier intake on the offspring’s intestinal compartment at weaning, as described **Fig. 3a**. Such an approach interestingly revealed that at weaning, and even in the absence of HFD or DSS insult, offspring from emulsifiers-exposed dams displayed increased body weight together with increased adipose deposition (**Fig. 3b-c**), as well as low-grade colonic inflammation evidenced by modestly but significantly elevated histopathological score (**Fig. 3d**). Next, to mechanistically investigate the effect of early-life microbiota alteration following maternal emulsifiers exposure on long-term disease susceptibility, we performed bulked RNA-seq on colonic samples from weanling pups (**Fig. 3e-h**). Such an approach revealed differentially expressed genes in CMC (23 genes, **Fig. 3e**) and P80 offspring (125 genes, **Fig. 3f**) compared to water control offspring. Gene enrichment pathways analysis revealed that pathways involved in response to microorganisms are highly impacted by maternal emulsifier consumption (**Fig. 3g, h**), aligning with the observed increased microbiota pro-inflammatory potential in CMC and P80 offspring (**Fig. 1g, i**). For example, CMC and P80 offspring were characterized by a decreased colonic expression of *Ifng*, encoding the INF-γ cytokine (**Fig. 3i**), for which microbiota-driven increased expression at weaning was previously reported to be central in driving long-term protective imprinting, a phenomenon termed weaning reaction ^1^. With the previous observation that intestinal immune cells are involved in such imprinting, we next assessed the intestinal immune compartment by investigating immune populations and cytokine *ex vivo* production in weanling mice, using mesenteric lymph node isolated cells. Such an approach revealed an increased expression of *Il1b* in mesenteric lymph nodes-derived cells after 24h *ex vivo* culture (**Fig. 3j**), aligning with the observed increased inflammatory tone in CMC and P80 offspring and highlighting the inflammatory skewing of intestinal immune cells activity. Quantification of immune cell populations using flow cytometry revealed that pups born from CMC-exposed dams displayed a significant decreased in B cells together with an increased in T cells (**Fig. 3k, l**). Moreover, despite no difference in overall T cell population in P80 offspring (**Fig. 3l**), we observed a reduction in total FoxP3^+^ T_reg_ (**Fig. 3m**), mostly driven by RorɣT^+^ tissue induced T_reg_ ^43^ rather than natural T_reg_ (**Fig. n, o**), suggesting that early life microbiota of P80 offspring is less efficient in establishing tolerant immune population.

**Figure 3.**
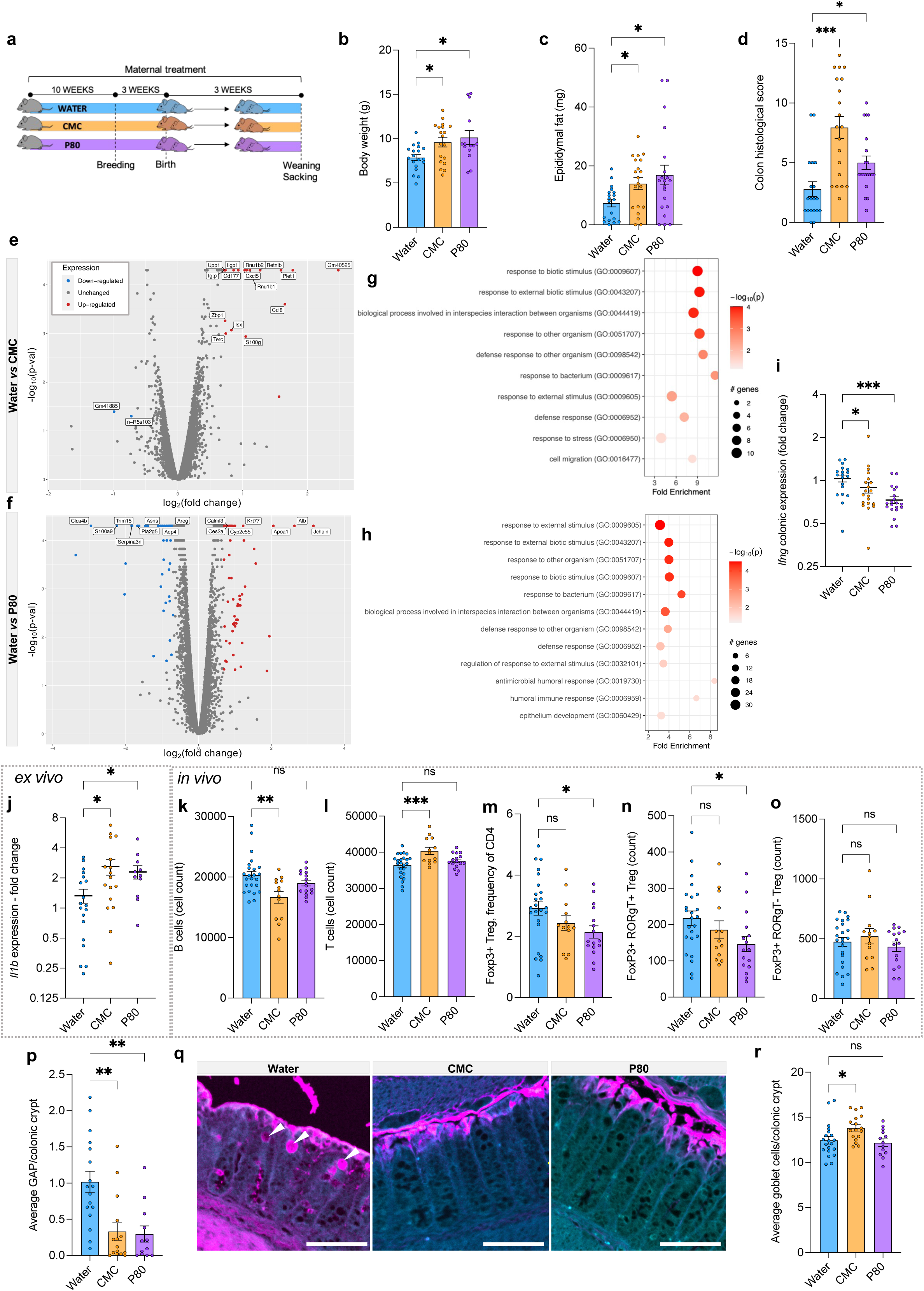
Maternal emulsifiers consumption induces intestinal inflammation and perturbation of the weaning reaction. Dams were subjected to 1% CMC or P80 in their drinking water for 10 weeks prior to breeding, and males offsprings were used just prior weaning for the analysis of intestinal inflammation and goblet-cell mediated molecular communication. (**a**) Schematic representation of the experimental design used. (**b**) Body weight, (**c**) periepididymal fat deposition and (**d**) histopathological scoring of HE stained colon samples at weaning. RNA seq was performed on colon samples collected at weaning; (**e-f**) volcano plots of genes expression in Water *vs* CMC offspring (**e**) or Water *vs* P80 offspring (**f**). (**g-h**) Significantly altered pathways between Water and CMC offspring (**g**) or between Water and P80 offspring (**h**). (**i**) Colonic expression fold change of *Ifng.* (**j**) *Il1b* fold change expression by mLN cells after 24h *ex vivo* culture. (**k-o**) Mesenteric lymph nodes (mLN) cells were analyzed by flow cytometry; (**k**) B220^+^ B cells, (**l**) CD3^+^ T cells, **(m)** Foxp3^+^ CD25^+^ CD127^-^ T_reg_, (**n**) Foxp3^+^ RORɣt^+^ T_reg_ and (**o**) Foxp3^+^ RORɣt^-^ T_reg_. (**p-q**) Quantification of colonic goblet-cell associated antigens passages at 18 days, expressed as the mean of GAP/crypt with the measurement of ovalbumin positive goblet cells performed in 50 colonic crypt per samples (**p**). Representative images of ovalbumin-labelled GAPs in transversal colonic sections (Ovalbumin labeled in pink, actin labeled in light blue, scale=50μm) (**q**). (**r**) Goblet cells per colonic crypt, quantified on alcian blue strained colonic section. Data are the means ± s.e.m (N = 13-24 from ≥ 3 independent litters). Significance was assessed by two-way ANOVA (**b-d, i-p, r**) and is indicated as follow: n.s., non-significant; * ≤ 0,05; **p ≤ 0,01; ***p ≤ 0,001.

With this observation of microbiota-driven impaired weaning reaction and breach in intestinal immune tolerance in emulsifier offspring, we next investigated early-life mechanism involved in driving *Ifng* expression and immune tolerance implementation. Among them, goblet cells associated passage (GAPs) are only observed during the early life window of opportunity and are suspected to play long lasting impact on intestinal health ^16^. Since colonic GAPs closure at weaning is under the control of MyD88 signaling ^16^, together with our observation that CMC and P80 offspring harbor microbiota with an increase expression of Microbe-Associated Molecular Patterns (MAMPs, **Fig. 1g**), we next quantified colonic GAPs in 18 day-old offspring from water-, CMC- or P80-treated dams. Using colonic instillation with fluorescently-labelled ovalbumin, followed by confocal-based quantification of ovalbumin positive goblet cells, we importantly observed a significant decrease in the number of colonic GAPs in CMC and P80 offspring (**Fig. 3p, q**), despite no reduction in the total number of goblet cells (**Fig. 3r**). Altogether, this suggest that maternal consumption of emulsifier drive an increased pro-inflammatory potential of the offspring microbiota, ultimately leading to premature GAPs closure during early life, hence preventing the establishment of proper host-microbiota interaction.

### Earl-life restoration of GAP opening fully prevents transgenerational long-lasting consequences of maternal emulsifier consumption

We next investigated whether such GAPs premature closure was directly involved in the observed deregulated inflammatory tone at weaning and the associated long-lasting consequences. For this purpose, pups from water-, CMC- or P80-treated dams were intraperitoneally injected daily from day 10 to day 21 with Tyrphostin, an inhibitor of the Epidermal Growth Factor Receptor (EGFR), known to prevent GAPs closure in the colon^16^ (**Fig. 4a**). As previously reported, such Tyrphostin treatment was sufficient to fully prevent GAPs reduction normally observed in CMC and P80 offspring (**Fig. 4b, c**). Such maintenance of GAPs opening fully prevented maternal emulsifier consumption-associated decreased in colonic expression of *Ifng* (**Fig. 4d**), suggesting restoration of a proper weaning reaction in these offspring. Of note, Tyrphostin treatment not only restored but increased GAPs opening, resulting in further increased *Ifng* expression (**Fig. S7a-f**) compared to control untreated offspring. This observation importantly suggest that GAPs-mediated antigen trafficking is involved in driving the increase *Ifng* expression reflecting the weaning reaction. Phenotypically, maintenance of GAPs opening in CMC and P80 offspring was sufficient to restore normal body weight at weaning (**Fig. 4e**), peri-epididymal fat deposition (**Fig. 4f**), as well as colonic inflammatory level (**Fig. 4g**).

**Figure 4.**
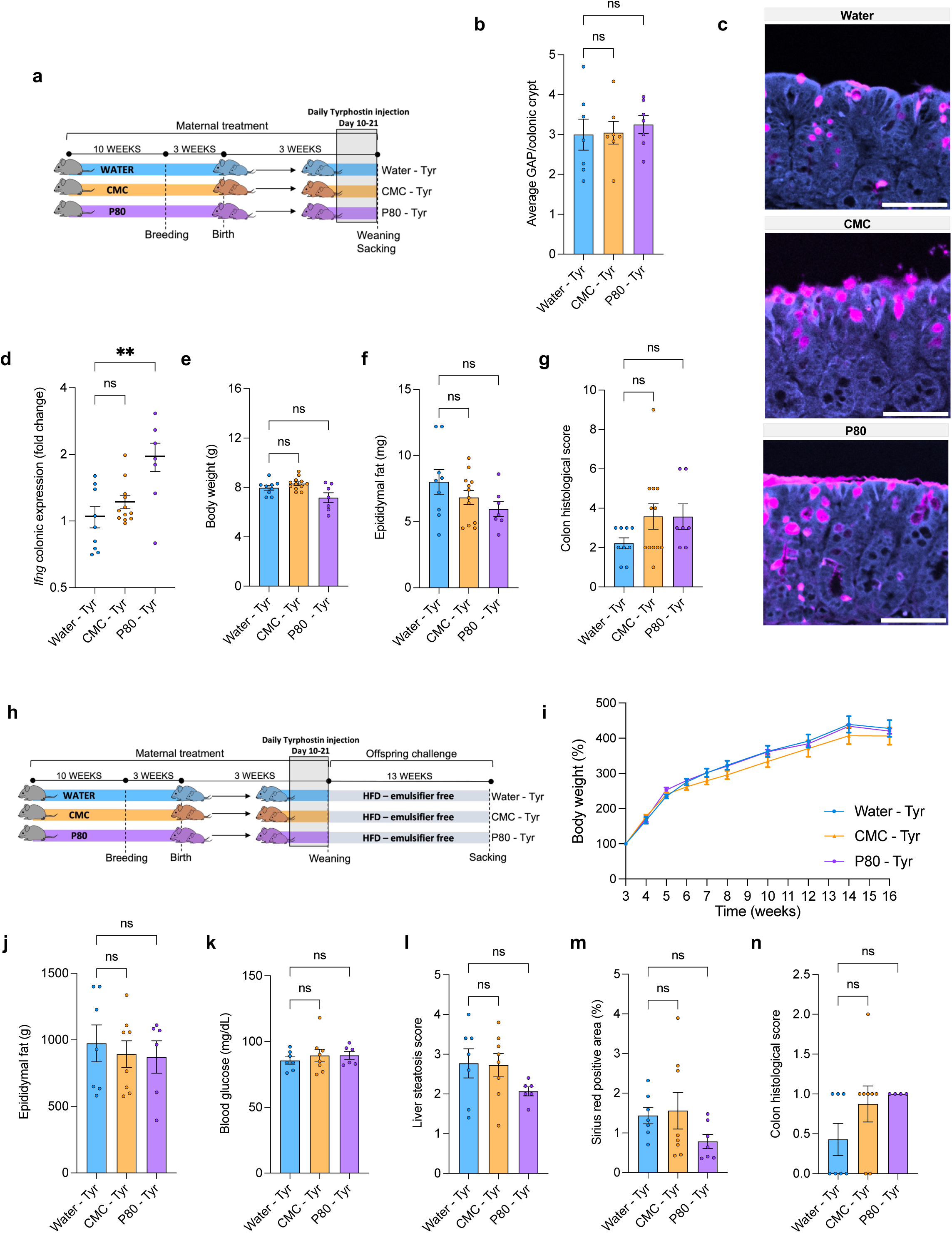
Chemically-induced GAP restoration fully prevents intestinal inflammation and susceptibility to diet-induced obesity normally associated with maternal intake of emulsifiers. Dams were subjected to 1% CMC or P80 in their drinking water for 10 weeks prior to breeding, and obtained pups were daily injected, from day 10 to day 21, with tyrphostin in order to maintain GAPs opening. Female offspring were sacked at weaning to quantify GAPs, intestinal inflammation and metabolism, while male offspring were weaned under emulsifiers-free conditions and challenged with a HFD for 13 weeks. (**a**) Schematic representation of the experimental design used. (**b)** Quantification of colonic goblet-cell associated antigens passages and (**c**) representative images of ovalbumin-labelled GAPs in transversal colonic sections (Ovalbumin labeled in pink, actin labeled in light blue, scale=50μm). (**d**) colonic expression fold change of *Ifng.* (**e**) Body weight at weaning, (**f**) periepididymal fat deposition and (**g**) histopathological scoring of HE stained colon samples. (**h**) Schematic representation of the experimental design used to investigate susceptibility to HFD-induced metabolic deregulations of tyrphostin-treated mice. (**i**) Body weight over time, with data being expressed as percentage compared to the body weight at weaning (week 3). (**j**) Peri-pididymal fat deposition measured upon sacking and (**k**) 15h fasting glycemia measured at week 14. (**l-m**) Liver histological samples were stained with HE or sirius red for steatosis scoring (**l**) and fibrosis quantification as the percentage of sirus red positive area (**m**). (**n**) Histopathological scoring of HE stained colon samples following HFD challenge. Data are the means ± s.e.m (N = 6-12 from ≥ 3 independent litters). Significance was assessed by one-way ANOVA (**b, d-g, j-n**) or two-way ANOVA (**i**) and is indicated as follow: n.s., non-significant; * ≤ 0,05; **p ≤ 0,01.

Finally, we investigated whether early life anticipated GAPs closure was also involved in long-lasting susceptibility to diet-induced obesity. Mice subjected to tyrphostin treatment between day 10 to day 21 were subsequently challenged with a HFD for 13 weeks, starting at weaning (**Fig. 4h**). Maintenance of GAPs opening in CMC-Tyrphostin and P80-Tyrphostin mice was sufficient to fully prevent the increased susceptibility to diet-induced metabolic deregulations normally observed in CMC and P80 offspring, as evidenced by normalization of body weight gain (**Fig. 4i**), peri-epididymal fat deposition (**Fig. 4j**), glycemia (**Fig. 4k**), liver steatosis and fibrosis (**Fig. 4l, m**), as well as intestinal inflammation (**Fig. 4n**). To conclude, these results suggest that the adverse transgenerational effects of maternal emulsifier consumption are driven by alterations in microbiota-host interaction through goblet cells associated passage.

## Discussion

Data have accumulated over the last 15 years, from our group and others, regarding the deleterious impact of dietary emulsifier on the intestinal microbiota, with subsequent detrimental consequences for host health. This includes the observation that dietary emulsifiers consumption is sufficient, in both mice models and human, to alter microbiota composition and increase its pro-inflammatory potential in a way that could drive intestinal inflammation and metabolic dysregulation ^26,28,29,37,38,44^. More recently, large epidemiological studies have reported positive associations between emulsifiers intake and risk of various chronic diseases ^30–32^. In these past research efforts, the potential trans-generational consequences of dietary emulsifier consumption were never investigated. While the role played by the early life microbiota in shaping long term host health has been recently documented, the exact actors at play, as well as the associated molecular mechanisms, remain largely unknow. Hence, we investigated here the potential consequence of maternal emulsifier consumption and its associated microbiota alterations on the next generation’s microbiota and health.

We observed that maternal intake of microbiota perturbator dietary emulsifiers was sufficient to induce offspring’s microbiota alteration in early-life, at both the compositional and functional levels. These alterations were associated with long-lasting susceptibility to diet-induced obesity and colitis, and cross-fostering approaches revealed a central role played by the intestinal microbiota in dictating such long-lasting consequences. We hypothesize that such long-lasting deleterious imprint of maternal-driven early-life microbiota alteration is mediated by impaired immune tolerance settlement in early-life, resulting in life-lasting hyper-inflammatory immune system. Indeed, in early-life, colonic goblet cells-associated antigen passages (GAPs) quantification led to the observation that offspring from emulsifier-exposed dams displayed a stark reduction in GAPs levels. Based on the previous observation that MyD88 signaling is central in modulating GAPs opening, with both caecal content from SPF mice and LPS luminal injection being sufficient to induce GAPs premature closure in the colon of WT but not in MyD88^-/-^ host ^16^, we are hypothesizing that microbiota encroachment within the normally sterile inner mucus layer contributes to premature MyD88 signaling prior weaning, thus driving colonic GAPs closure. This is further strengthened by our observation that maternal emulsifier consumption drives an increased level of bioactive flagellin in offspring, which we believe contributes to activate MyD88-dependant signaling, especially during this early-life time period characterized by high *Tlr5* colonic expression ^3,45^. Importantly, chemical maintenance of GAPs opening was sufficient to fully prevent long lasting transgenerational consequences of maternal emulsifier consumption, highlighting the crucial role played by GAPs-mediated early-life host-microbiota communication in host long-lasting health, as illustrated **Fig. S8**. Furthermore, increased GAP levels was strongly associated with IFN-γ levels, suggesting that GAP modulate the previously-described IFN-γ pick observed during the weaning reaction (**Fig. S7, S8**). Increased IFN-γ following Tyrphostin-induced GAPs opening boost was even further enhanced in CMC and P80 offspring compared to water offspring, which we suspect to be driven by the increased immunostimulatory potent of the luminal microbiota-derived antigens trafficking through GAPs in these emulsifier-treated mice.

We observed that microbiota composition differed between CMC- and P80-exposed dams, as previously described ^26,37^, suggesting compound-specific effects. Additionally, P80- and CMC-specific metabolic deregulations were noted in the offspring following a HFD regimen, suggesting that the presence of select microbiota members during early life may lead to distinct host imprinting. In addition, in CMC offspring, some phenotypes were not fully restored upon microbiota normalization at birth, suggesting that this compound and/or the associated altered maternal microbiota may have an *in utero* impact on the offspring. During fetal life, mother’s microbiota could indeed produce compounds that are transferred to the fetus in a way that can impact innate immune cell populations ^46^. Another point in dire need of further investigations relates to the role played by the recently observed heterogeneous response to emulsifiers in humans on the transgenerational effects described here. We indeed recently reported that microbiota inter-individual variation are key in driving host responses to emulsifier consumption^28^, leading to some individuals being highly sensitive to emulsifier-induced perturbations. Investigating the trans-generational consequences of such personalized responsiveness toward emulsifier appears essential to fully appreciate the impact of maternal emulsifier consumption across generations.

To conclude, our observation that maternal emulsifiers intake induces offsprings microbiota alteration in a way that disrupt host-microbiota interactions, leading to long-lasting susceptibility to metabolic and inflammatory diseases, underscores the need to carefully consider maternal exposure and dietary recommendations to ensure the long-term health of future generations.

## Methods

### Mice

Four-week-old male and female C57Bl/6J mice were housed in cages of five individuals and maintained under a 12-hour light/dark cycle. The male mice were housed with ad libitum access to standard chow diet and water, while the female mice were divided into three groups: a water-treated group, a group treated with 1% carboxymethyl cellulose (CMC), and a group treated with 1% polysorbate 80 (P80) in the drinking water. Cages for all groups were changed every other week. After 10 weeks of treatment, breeding pairs were formed by pairing two females with one male. Pregnant females were subsequently individually housed from gestation until weaning. On the day of birth, pups were either left with their biological mother or cross-fostered to dams with age-matching and size-matching litters. The resulting pups were categorized as follows:

#### Experiment 1 – Fig. 1, 2, S2, S4

Pups born to water-, CMC-, or P80-treated dams were either kept by their biological mother or cross-fostered with water-treated dams at birth. Male pups were weaned at 21 days-old under emulsifier-free conditions, with ad libitum access to water, and then subjected to a high-fat diet (HFD) for 13 weeks.

#### Experiment 2 – Fig. 2, S5, S6

Pups born to water-, CMC-, or P80-treated dams were either kept by their biological mother or cross-fostered with water-, CMC-, or P80-treated dams at birth. Male pups were weaned at 21 days old under emulsifier-free conditions, with ad libitum access to standard chow diet and water. After 4 weeks, these mice were subjected to 1% dextran sodium sulfate (DSS) in the drinking water for 7 days.

#### Experiment 3 – Fig. 3

Half of the male pups born to water-, CMC-, or P80-treated dams were used at day 18 to quantify goblet cells-associated antigen passage (GAP), while the other half were euthanized on the day of weaning for intestinal inflammation quantification.

#### Experiment 4 – Fig. 4

Pups born to water-, CMC-, or P80-treated dams were injected daily intraperitoneally (IP) with 0.5 μg of Tyrphostin per gram of body weight from day 10 to day 21 ^16^. Female pups were used to quantify GAP at weaning. Male pups were weaned at 21 days old under emulsifier-free conditions, with ad libitum access to water, and then subjected to a HFD for 13 weeks. Fasting glycemia was measured after 10 weeks of HFD.

Body weight was measured and feces were collected longitudinally throughout the experiments. Upon euthanasia, mice were anesthetized with isoflurane, blood samples collected, and mice were euthanized by cervical dislocation. Measurements of colon length, colon weight, spleen weight, and adipose weight were recorded, and tissue samples were collected for downstream analysis. Animal welfare and experimental protocols followed the ARRIVE guidelines (Animal Research: Reporting of *in vivo* Experiments). All procedures involving animals were approved by the French Ministère de la l’enseignement supérieur, de la recherche et de l’innovation, APAFIS#24788-2019102806256593 v8.

### Milk collection

Following 6 hours of pups isolation from their mother, dams are injected IP with 1.5UI of Ocytocine-S, (Sigma, O3251) prior anesthesia with isoflurane for milk collection. Milk samples were snap frozen for downstream analysis.

### Colonic GAP staining and quantification

Pups were subjected to intro-colonic injection with 0.25mg of Ovalbumin Alexa Fluor 647 conjugated (Invitrogen 034784). After 1h, mice were euthanized and colons were opened longitudinally and washed twice in PBS prior fixation in 4% paraformaldehyde. Following embedding in paraffin with a vertical orientation, five-μm sections were cut and dewaxed by bathing in xylene at 60 °C for 10 min, xylene at room temperature for 10 min, and 99.5% ethanol for 10 min. Sections were marked using a PAP pen (Sigma, St. Louis, MO, USA) and block solution (5% FBS in PBS) was added for 30 min at 4 °C. Mucin-2 primary antibody (rabbit H-300, [C3], C-term, Genetex, GTX100664) was diluted to 1:100 in block solution and applied overnight at 4 °C. After washing 3 × 10 min in PBS, block solution containing anti-rabbit Alexa 488 secondary antibody diluted to 1:300, PhalloidinTetramethylrhodamine B isothiocyanate (SigmaAldrich) at 1 mg ml−1 and Hoechst 33258 (Sigma-Aldrich) at 10 mg ml−1 was applied to the section for 2 h. After washing 3 × 10 min in PBS slides were mounted using Prolong anti-fade mounting media (Life Technologies) and kept in the dark at 4 °C. Observations and measurement of the distance between bacteria and epithelial cell monolayer were performed with a Spinning Disk IXplore using the Olympus cellSens imaging software 421 (V2.3) at a frame size of 2048 × 2048 with 16-bit depth. A 405 nm laser was used to excite the 422 Hoechst stain (epithelial DNA), 488 nm for Alexa Fluor 488 (mucus), 488 nm for Phalloidin (actin), 423 and 640 nm for Alexa Fluor 647 (ovalbumin). Samples were imaged with a ×10 objective, and GAP identified as ovalbumin-labeled goblet cells were quantified over 50 villus using QuPath 0.5.0 software.

### Fasting blood glucose measurement

After 10 weeks of HFD, mice were placed in a clean cage and fasted for 15 hours. Blood glucose concentration was then determined using a Nova Max Plus Glucose Metre and expressed in mg/ dL.

### Steatosis and fibrosis scoring

Following euthanasia, livers were harvested and fixed in 4% paraformaldehyde, embedded in paraffin, five-μm sectioned, and stained with hematoxylin and eosin (steatosis) or Sirius red (fibrosis). Steatosis was evaluated blindly and as previously described ^47^. Briefly, on 5 equal size section of each slide, a score between 0 and 3 was given to macrovesicular steatosis, microvesicular steatosis, hypertrophy and number of inflammation foci ^47^. Fibrosis was determined by measuring the percentage of sirius positive area on the whole section.

### Staining of colonic tissue and histopathologic analysis

Following euthanasia, colons (proximal colon, 2 first cm from the cecum) were placed in Carnoy’s fixative solution (60% methanol, 30% chloroform, 10% glacial acetic acid). Tissues were then washed in methanol 2 × 30 min, ethanol 2 × 15 min, ethanol/ xylene (1:1) 15 min, and xylene 2 × 15 min, followed by embedding in paraffin with a vertical orientation. Tissues were sectioned at 5-μm thickness and stained with hematoxylin & eosin (H&E) using standard protocols. H&E-stained slides were assigned four scores based on the degree of epithelial damage and inflammatory infiltrate in the mucosa, submucosa, and muscularis/serosa. Each of the four scores was multiplied by 1 if the change was focal, 2 if it was patchy, and 3 if it was diffuse, as previously described^48^. The four individual scores per colon were added, resulting in a total scoring range of 0–36 per mouse.

Colonic sections (4 μm) were also stained with Alcian Blue, preferentially staining mucopolysaccharides, and 17–23 crypts of 3 regions per colonic sections were randomly selected per animal to determine goblet cell number per crypt.

### Liver and colon mRNAs extraction and q-RT-PCR analysis

Distal colons were collected during euthanasia and placed in RNAlater. Total mRNAs were isolated from colonic tissues using TRIzol (Invitrogen, Carlsbad, CA) according to the manufacturer’s instructions and as previously described^48^. Quantitative RT-PCR was performed using the Qiagen kit QuantiFast® SYBR® Green RTPCR in a LigthCycler® 480 instrument (Roche Molecular Systems, Inc.) with mouse gene-specific oligonucleotides (Supplementary Table 1). Gene expressions are presented as relative values using the Ct approach with the 36B4 housekeeping gene.

### Ex-vivo cell culture and cytokine quantification

Cells from mesenteric lymph nodes (mLN) were isolated as follow: mLN were collected in HBSS prior cell dissociation by pushing cells through a 100μm filter with a syringe plug. Cells were then washed once in HBSS before resuspending 500,000 cells in RPMI for culture. After 24 hours, cells are collected and mRNAs were extracted as described above.

### Small intestine, colon and mLN cell preparation, and flow cytometry analysis

mLN cells were isolated as previously described in the method section. For small intestinal and colonic lamina propria cells isolation, Peyer’s Patches were removed, and whole small intestine and colon were opened longitudinally, cut into pieces and incubated for 30 minutes in 30mM EDTA, washed extensively, and incubated for 15 min at least twice in 1mg/mL collagenase D and 1U/mL DNAse I in HBSS. These preparations were then pushed through a 100μm filter to generate single cell suspensions. Cells were separated by a 40/80% (w/v) Percoll (GE Healthcare) density gradient and washed prior to staining for flow cytometry analysis. Cells were then pre-incubated with Zombie UV™ Fixable Viability Kit for 30 minutes, Fc-Block for 15 minutes, and stained for 20 minutes with the following antibodies: anti-CD45 PERCP (530-F11; Biolegend), anti-B220 Pacific Blue (RA3-6B2; BD Biosciences), anti-CD3 BV711 (17A2; Biolegend). Cells isolated from ileum and mLN were additionally stained with anti-CD138 BB515 (281-2; BD Biosciences), anti-IgA BV786 (C10-1; BD Biosciences), anti-CD4 (GK1.5; BD Biosciences), anti-CD8 V500 (53-6.7; BD Biosciences), anti-CD127 BV605 (A7R34; Biolegend), anti-CD25 BV650 (PC61; BD Biosciences) and intracellular staining was performed following fixation and permeabilization with anti-Tbet PE-CF594 (O4-46; BD Biosciences), anti-Gata3 AF700 (TWAJ; eBioscience), anti-RorɣT PE (Q31-378; BD Biosciences), anti-FoxP3 APC (FJK16s; eBioscience). Cells isolated from colon were additionally stained with anti-MHCII APC-Cy7 (M5/114.15.2; Biolegend), anti-CD115 APC (AFS98; eBioscience), anti-Ly6G/C BV650 (RB6-8C5; Biolegend), anti-CD103 PE-CF594 (M290; BD Biosciences), anti-CD11c BB515 (NA18; BD Biosciences), anti-CX3CR1 AF700 (SA011F11; Biolegend), anti-CD206 PE (C068C2; Biolegend), anti-CD27 BV786 (LG3A10; BD Biosciences) and anti-CD11b BV421 (M1/70; BD Biosciences). Samples were acquired on a BD LSRFortessa™ Cell Analyzer, and data were analyzed using a non-supervised approach using the Omiq software.

### Colonic mRNAs sequencing

Distal colon was collected during euthanasia and placed in RNAlater. Total mRNAs were isolated from colonic tissues using TRIzol (Invitrogen, Carlsbad, CA) according to the manufacturer’s instructions and as previously described^48^.

#### Library preparation and sequencing

cDNA library was prepared using the Invitrogen^TM^ Collibri^TM^ Stranded RNA library Prep Kit for Illumina^TM^ with Collibri^TM^ H/M/R rRNA Depletion Kit according to the manufacturer’s instructions and starting with 500ng of purified RNAs. Briefly, rRNA were first depleted, and enriched mRNAs subsequently used for fragmentation, adaptors ligation and reverse transcription. After purification, libraries were PCR-enriched, further purified, and quantified and quality-assessed on an Agilent^TM^ 2100 Bioanalyzer^TM^ instrument. A master library was generated from the purified products in equimolar ratios. The pooled product was quantified using Qubit and sequenced using an Illumina Next-Seq sequencer (paired-end reads, 2x750 bp).

#### Data analysis

Cutadapt online tool ^49^ was used to remove adapter sequences as well as trim sequences from the first low quality (<28) base. High quality reads longer than 20 nucleotides were then aligned to the mm10 *mus musculus* reference genome using Bowtie2^50^. Gene expression levels were next measured using Cufflinks^51^ and differentially expressed genes between conditions were identified using Cuffdiff^51^. Fragments Per Kilobase of transcript per Million mapped reads (FPKM) unit was used and Log2 fold changes and p-values were computed for each comparison of interest. Gene level volcano plots were generated through R (R version 4.2.3) and differentially expressed genes enrichment analysis was performed using the edgeR R package^52^ (package version 3.40.2) to identify genes with Log2 fold changes ≥ log(2) or ≤ -log(2) and p-value ≤ 0.05. Differentially expressed genes pathways enrichment analysis was performed using the PANTHER classification sytem^53^ and significant pathways with FDR ≤ 0.05 and more than 6 DEG involved were represented as enrichment plot (enrichment_chart, R package v2.3.0). Unprocessed sequencing data are deposited in the European Nucleotide Archive under accession number XXXXXXX.

### Caecal metabolites quantification

Proton nuclear magnetic resonance (^1^H NMR) – based metabolomics was performed on 100 mg of frozen caecal content as previously described ^54^. Data pretreatment and statistical analysis with principal component analysis was performed using full resolution spectra in MATLAB (R2021a; MathsWorks, Inc.).

### Maternal milk metabolites qualification and detection of CMC and P80

Maternal milk metabolites were quantified using ^1^H NMR-based metabolomics as described in our previous study with minor modifications ^55^. Approximately 200 μl of milk were mixed 1 mL of methanol:chloroform (2:1 v/v), vortexed for 30 sec, and then 133 µL of nanopure distilled water and 333ul of chloroform were sequentially added to the sample, vortexed for 30 s, and kept overnight at 4°C. After centrifugation at 6,000 g at 4°C for 20 min, the polar phase was collected in a 2 mL microcentrifuge tube and dried down using a SpeedVac vacuum concentrator. The dried extract was resuspended in 0.55ml of 0.1 M PBS (50% D2O, 0.005% w/v TSP), and after centrifugation at 17000 g at 4°C for 10 min, the supernatant was transferred into 5mm NMR tube for NMR analysis. The 1D ^1^H spectra of milk extracts were acquired at 298 K using a Bruker Avance NEO 600 MHz spectrometer equipped with a SampleJet sample changer (Bruker Biospin, Germany). The standard pulse sequence (noesygppr1d) was used for recording ^1^H NMR experiments with pre-saturation water suppression during relaxation and mixing time. All spectra were processed automatically with Chenomx NMR Suite 10 (Chenomx Inc, Edmonton, Alberta, Canada), and each spectrum was checked and adjusted manually for phase and baseline to satisfy quality requirements. Metabolites were identified and quantified using the built-in metabolite library and fitting algorithm in the Chenomx software, combined with the known concentrations internal standard (TSP, 0.29 mM).

### Quantification of fecal IgA-coated bacteria

IgA-coated bacteria were quantified as previously described ^56^. In brief, frozen fecal samples were thoroughly homogenized in PBS to a final concentration of 20 mg/mL. Fecal suspensions were filtered through a 40-μm sterile nylon mesh, then centrifuged at 50 × g, for 15 min at 4 °C. 200 μL of supernatant was then washed with 1 mL PBS and centrifuged at 8,000 × g, for 5 min at 4 °C. Resulting bacterial pellets were resuspended in 100 μl blocking buffer (staining buffer containing 20% Normal Rat Serum) and incubated for 20 min on ice before being stained with 100 μl of staining buffer containing PE-conjugated Anti-Mouse IgA (mA-6E1; eBioscience) for 30 min on ice, in the dark. Following two washes with staining buffer, pellets were resuspended in 200 μL of FACS buffer (PBS, 1% Normal Rat Serum). Data acquisition was performed on a Beckman Coulter Gallios flow cytometer. For each sample, 100,000 events were recorded and data was analyzed using FlowJo software v.10.8.2.

### Milk total IgA and anti-flagellin/LPS IgA quantification by ELISA

Milk samples were resuspended 1 in 10 in collection media consisting of 0.05 mg soybean trypsin inhibitor per ml of a 3:1 mixture of 1× PBS and 0.1 M EDTA, pH 7.4. Following centrifugation at 1800 rpm for 10 min, the supernatant was centrifuged again at 14,000 rpm for 15 min at 4 °C, and final supernatant was collected and stored with 20% glycerol and 2 mM phenylmethylsulfonyl fluoride. Quantification of total IgA, anti-flagellin and anti-LPS IgA was performed by coating 96-well microtiter plates (Costar, Corning, New York) with goat anti-mouse IgA (Southern Biotech), or 100 ng/well of laboratory made Salmonella Typhimurium-derived flagellin, or 2μg/well lipopolysaccharides (from E. coli 0128: B12, Sigma) in 9.6 pH bicarbonate buffer overnight at 4 °C. Serum or fecal samples from mice were then applied either pure at a final dilution of 1 in 10 000 for total IgA ELISA or 1 in 50 for anti-flagellin/LPS IgA ELISA for 1 h at 37 °C. After incubation and washing, the wells were incubated with horseradish peroxidase linked anti-mouse IgA (Southern Biotech, 1040–05). Quantification of immunoglobulin was then developed by the addition of 3,3′,5,5′-Tetramethylbenzidine and the optical density was calculated by the difference between readings at 450 nm and 540 nm.

### Microbiota analysis by 16 S rRNA gene sequencing

16S rRNA gene amplification and sequencing were done using the Illumina MiSeq technology following the protocol of Earth Microbiome Project with their modifications to the MOBIO PowerSoil DNA Isolation Kit procedure for extracting DNA (www.earthmicrobiome.org/emp-standard-protocols). Bulk DNA were extracted from frozen extruded feces using a PowerSoil-htp kit from MoBio Laboratories (Carlsbad, California, USA) with mechanical disruption (bead-beating). The 16S rRNA genes, region V4, were PCR amplified from each sample using a composite forward primer and a reverse primer containing a unique 12-base barcode, designed using the Golay error-correcting scheme, which was used to tag PCR products from respective samples ^57^. We used the forward primer 515F 5’-*AATGATACGGCGACCACCGAGATCTACACGCT*XXXXXXXXXXXX**TATGGTAATT*GT*** <u>GTGYCAGCMGCCGCGGTAA</u>-3’: the italicized sequence is the 5’ Illumina adapter, the 12 X sequence is the golay barcode, the bold sequence is the primer pad, the italicized and bold sequence is the primer linker and the underlined sequence is the conserved bacterial primer 515F. The reverse primer 806R used was 5’-*CAAGCAGAAGACGGCATACGAGAT***AGTCAGCCAG*CC***<u>GGACTACNVGGGTWTCTAAT</u>-3’: the italicized sequence is the 3’ reverse complement sequence of Illumina adapter, the bold sequence is the primer pad, the italicized and bold sequence is the primer linker and the underlined sequence is the conserved bacterial primer 806R. PCR reactions consisted of Hot Master PCR mix (Quantabio, Beverly, MA, USA), 0.2 μM of each primer, 10–100 ng template, and reaction conditions were 3 min at 95 °C, followed by 30 cycles of 45 s at 95 °C, 60 s at 50 °C, and 90 s at 72 °C on a Biorad thermocycler. Products were then visualized by gel electrophoresis and quantified using Quant-iT PicoGreen dsDNA assay (Clariostar Fluorescence Spectrophotometer). A master DNA pool was generated in equimolar ratios, subsequently purified with Ampure magnetic purification beads (Agencourt, Brea, CA, USA) and sequenced using an Illumina MiSeq sequencer (paired-end reads, 2 × 250 bp) at the Genom’IC platform (INSERM U1016, Paris, France). Unprocessed sequencing data are deposited in the European Nucleotide Archive under accession number XXXXXXX.

### 16S rRNA gene sequence analysis

16S rRNA sequences were analyzed using QIIME2—version 2022 ^58^. Sequences were demultiplexed and quality filtered using the Dada2 method ^59^ with QIIME2 default parameters in order to detect and correct Illumina amplicon sequence data, and a table of QIIME2 artifact was generated using the folowing dada2 command: qiime dada2 denoise-paired --i-demultiplexed-seqs demux.qza --p-trim-left-f 0 --p-trim-left-r 0 --p-trunc-len-f 180 --p-trunc- len-r 180 --o-representative-sequences rep-seqs-dada2.qza --o-table table-dada2.qza --o- denoising-stats stats-dada2.qza --p-n-threads 6. A tree was next generated, using the align-to- tree- mafft-fasttree command, for phylogenetic diversity analyses, and alpha and beta diversity analyses were computed using the core-metrics-phylogenetic command. Principal coordinate analysis (PCoA) plots were used to assess the variation between the experimental group (beta diversity). For taxonomy analysis, features were assigned to operational taxonomic units (OTUs) with a 99% threshold of pairwise identity to the Greengenes reference database (version Greengenes2 2022.10).

### Identification of microbiota members significantly altered in their relative abundance

Microbiota members presenting significant changes in their abundance between comparison of interest were identified using MaAsLin2 (Microbiome Multivariable Associations with Linear Models, version 2)^60^. MaAsLin2 analysis (R version 4.1.2, Maaslin2 version 1.12.0 package) was conducted using relative abundance data for microbiota members identified at the species level. Microbiota members were reported as significantly altered in their relative abundance if corrected *q*-value < 0.05 (**Fig. 1e**) or *p*-value < 0.05 (**Fig. S6f, i**) .

### Identification of bacteria targeted by maternal milk IgA

Milk samples were initially diluted at a ratio of 1:50 with a bacterial suspension derived from RagKO mice fecal material^61^, which was previously resuspended in phosphate-buffered saline (PBS) to achieve a final concentration of 100 mg/mL after homogenization and filtration. Following a one-hour incubation period at 37°C, IgA-coated bacteria were identified by flow cytometry following IgA staining, and a population enriched with IgA-positive bacteria, comprising 100,000 events, was isolated using a S3e Cell Sorter Biorad. Subsequently, DNA extraction was carried out from the sorted sample containing IgA-coated bacteria, as well as from the mixture resulting from the incubation of milk in the fecal suspension from Rag mice fecal material. The QIAamp Fast DNA Stool Mini Kit (Qiagen) was next used for DNA extraction. For taxonomic profiling, 16S sequencing was conducted, enabling the identification of bacteria bound by IgA. Significantly enriched bacteria were determined by comparing the IgA-coated bacteria enriched sorted sample with the product obtained from the incubation of milk in the fecal suspension from Rag mice fecal material, utilizing the MaAsLin2 approach.

### Bioactive flagellin and LPS fecal load quantification

Levels of fecal bioactive flagellin and LPS were quantified, as previously described ^36^, using human embryonic kidney (HEK)-Blue-mTLR5 and HEK-Blue-mTLR4, respectively (Invivogen, San Diego, California). Fecal material were resuspended in PBS to a final concentration of 100 mg.mL^-1^ and homogenized for 15 minutes using a vortex. We then centrifuged the samples at 8000g for 15min and serially diluted the resulting supernatant and applied to mammalian cells. Purified *Escherichia coli* flagellin and LPS (Sigma, St Louis, Missouri) were used for standard curve determination using HEK-Blue-mTLR5 and HEK-Blue-mTLR4, respectively. After 24 h of stimulation, we applied cell culture supernatant to QUANTI-Blue medium (Invivogen, San Diego, California) and measured alkaline phosphatase activity at 620 nm after 30 min.

### AhR ligands quantification in milk samples

Levels of AhR ligands were measured in milk samples using HT-29-Lucia™ AhR reporter cells (Invivogen, San Diego, California). Briefly, milk samples diluted 1:5 in PBS were centrifuged at 8,000g, and the supernatant was applied to HT-29-Lucia™ AhR reporter cells. Luminescence was immediately measured, and sample concentrations were calculated based on the luminescence measurements of HT-29-Lucia™ AhR reporter cells stimulated with a known amount of the AhR ligand FICZ.

### Immunostaining of mucins and localization of bacteria by fluorescent in situ hybridization

Mucus immunostaining was paired with fluorescent *in situ* hybridization (FISH), as previously described ^26,62^, in order to analyze bacteria localization at the surface of the intestinal mucosa. In brief, colonic tissues (proximal colon, second cm from the caecum) containing fecal material were placed in methanol-Carnoy’s fixative solution (60% methanol, 30% chloroform, 10% glacial acetic acid) for a minimum of 3 h at room temperature. Tissues were then washed in methanol 2x30 min, ethanol 2x15 min, ethanol/xylene (1:1) 15 min and xylene 2x15 min, followed by embedding in paraffin with a vertical orientation. Five-μm sections were cut and dewaxed by preheating at 60°C for 10 min, followed by bathing in xylene at 60°C for 10 min, xylene at room temperature for 10 min and 99.5% ethanol for 10 min. The hybridization step was performed at 50°C overnight with an EUB338 probe (59-GCTGCCTCCCGTAGGAGT- 39, with a 59 Alexa 647 label) diluted to a final concentration of 10 mg.mL^-1^ in hybridization buffer (20 mM TrisHCl, pH 7.4, 0.9 M NaCl, 0.1% SDS, 20% formamide). After washing for 10 min in wash buffer (20 mM Tris-HCl, pH 7.4, 0.9 M NaCl) and 3x10 min in PBS, a PAP pen (Sigma, St. Louis, Missouri) was used to mark around the section and block solution (5% FBS in PBS) was added for 30 min at 4°C. Mucin-2 primary antibody (rabbit H-300, [C3], C-term, Genetex, GTX100664) was diluted to 1:100 in block solution and applied overnight at 4°C. After washing 3x10 min in PBS, block solution containing anti-rabbit Alexa 488 secondary antibody diluted to 1:300, PhalloidinTetramethylrhodamine B isothiocyanate (Sigma-Aldrich) at 1 mg.ml^-1^ and Hoechst 33258 (Sigma-Aldrich) at 10 mg.ml^-1^ was applied to the section for 2 h. After washing 3x10 min in PBS slides were mounted using Prolong anti- fade mounting media (Life Technologies) and kept in the dark at 4°C. Observations and measurement of the distance between bacteria and epithelial cell monolayer were performed with a Spinning Disk IXplore using the Olympus cellSens imaging software 421 (V2.3) at a frame size of 2,048 x 2,048 with 16-bit depth. A 405nm laser was used to excite the 422 Hoechst stain (epithelial DNA), 488nm for Alexa Fluor 488 (mucus), 488nm for Phalloidin (actin), 423 and 640nm for Alexa Fluor 647 (bacteria). Samples were imaged with a 20x objective.

### Maternal treatment prediction based on offspring microbiota composition

The association between offspring microbiota composition and maternal treatment was assessed trough the prediction of maternal treatment based on offspring’s microbiota composition data. Prediction of CMC treatment (outcome CMC or WATER, **Fig. S2a-d**), P80 treatment (outcome P80 or WATER, **Fig. S2e-h**) was performed by computing receiver operating characteristic (ROC) curves (R version 4.1.2, randomForest 4.7-1.1 package, ROCR package) using training data set and validation data set containing randomly affected 80% and 20% of mice, respectively. Data set contained relative abundance data for microbiota members identified at the species level at weeks 4, 8, 12 and 16 of age. ROC calculation was repeated 50 times with random sampling of the training and validation data and area under curve (AUC) measurement for each iteration. Mean AUC and standard deviation are presented for each graph.

### Statistical analysis

Significance was determined using, when normality and homoscedasticity postulates were valid, one-way group analysis of variance (ANOVA) with Sidak’s multiple comparisons test. Significance of data that did not respect normality and homoscedasticity postulates was tested using Kruskal-Wallis corrected for multiple comparisons with a Dunn’s test or Brown-Forsythe and Welch ANOVA corrected for multiple comparisons with a Dunnett test, respectively. Significance of longitudinally-measured data was assessed using two-way ANOVA corrected for multiple comparisons with a Sidak’s test. Clustering significance in PCoA plot was determined using a ANOSIM analysis of similarities test. Differences were noted as significant *p ≤ 0,05; ** p ≤ 0,01; *** p ≤ 0,001; **** p ≤ 0,0001; n.s. indicates nonsignificant.

## Funding

This work was supported by a Starting Grant from the European Research Council (ERC) under the European Union’s Horizon 2020 research and innovation program (grant agreement No. ERC-2018-StG- 804135), ANR grants EMULBIONT (ANR-21-CE15-0042-01) and DREAM (ANR-20-PAMR-0002), DIM BioConvergence for Health (BioConvS) and Région Ile-de-France for the founding the cell sorted used in this study, and the national program “Microbiote” from INSERM. AP and FH are supported by the USDA National Institute of Food and Agriculture and Hatch Appropriations under #PEN4917 and Accession #7006412. CD is supported by a fellowship from the *Fondation pour la Recherche Médicale* (FRM). No funders had any role in the design of the study and data collection, analysis, and interpretation, nor in manuscript writing.

## Acknowledgment

The authors thank the Hist’IM and the Genom’IC platforms (INSERM U1016, Paris, France) for their help. The authors thank Enzo Manchon (INSERM, Saint Louis hospital, Paris France) for his constructive help with flow cytometry analysis and the use of Omiq license.

---

**Figure S1.**
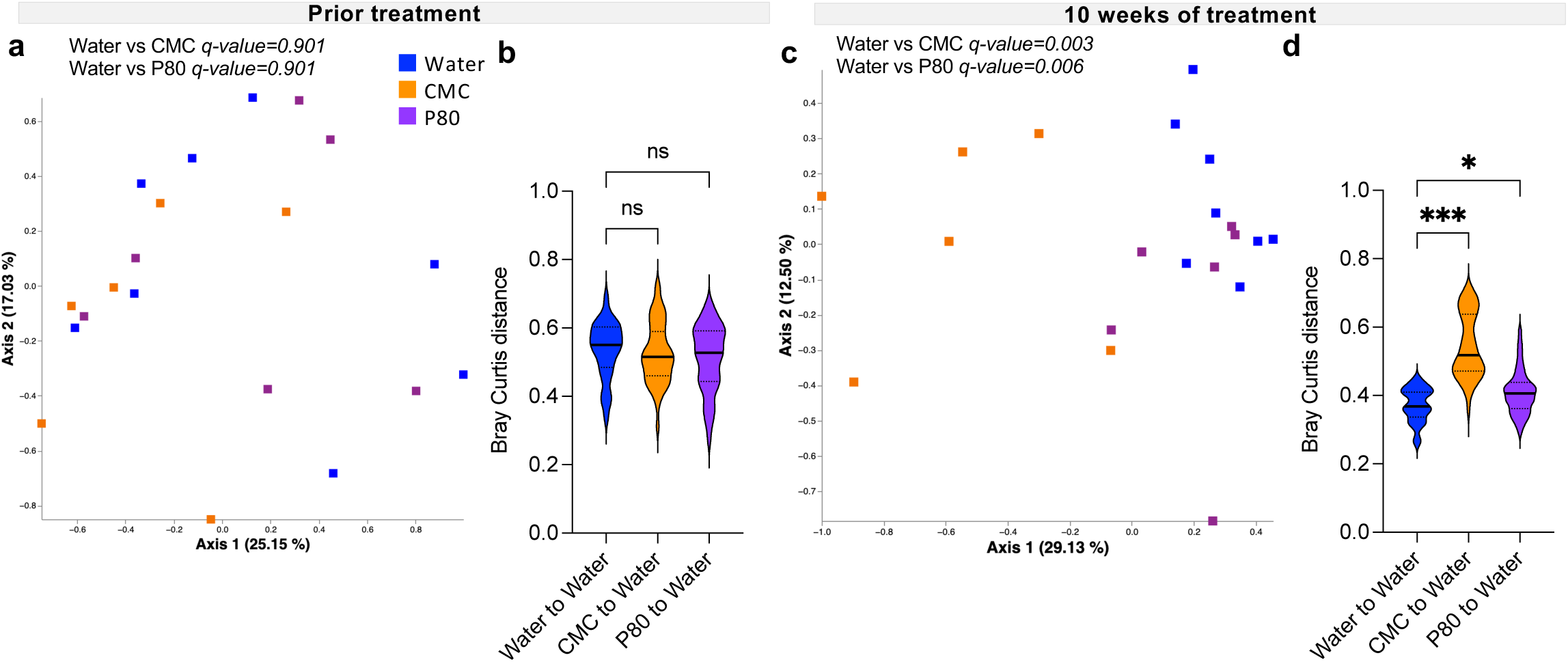
Chronic exposure to dietary emulsifiers impacts dam’s intestinal microbiota composition. Dams were subjected to 1% CMC or P80 in their drinking water for 10 weeks. (**a-b**) Dam’s microbiota composition prior treatment, represented using principal coordinates analysis of the Bray–Curtis distance (**a**) or using Bray Curtis distance separating the various experimental groups from the water-treated group (**b**). (**c-d**) Dam’s microbiota composition after 10 weeks of treatment, represented using principal coordinates analysis of the Bray–Curtis distance (**c**) or using Bray Curtis distance separating the various experimental groups from the water-treated group (**d**). N = 6-8 from ≥ 3 independent cages. Significance was assessed by ANOSIM (**a, c**), or one-way ANOVA (**b, d**), and is indicated as follow: n.s., non- significant; * p ≤ 0.05; ** p ≤ 0.01; *** p ≤ 0.001.

**Figure S2.**
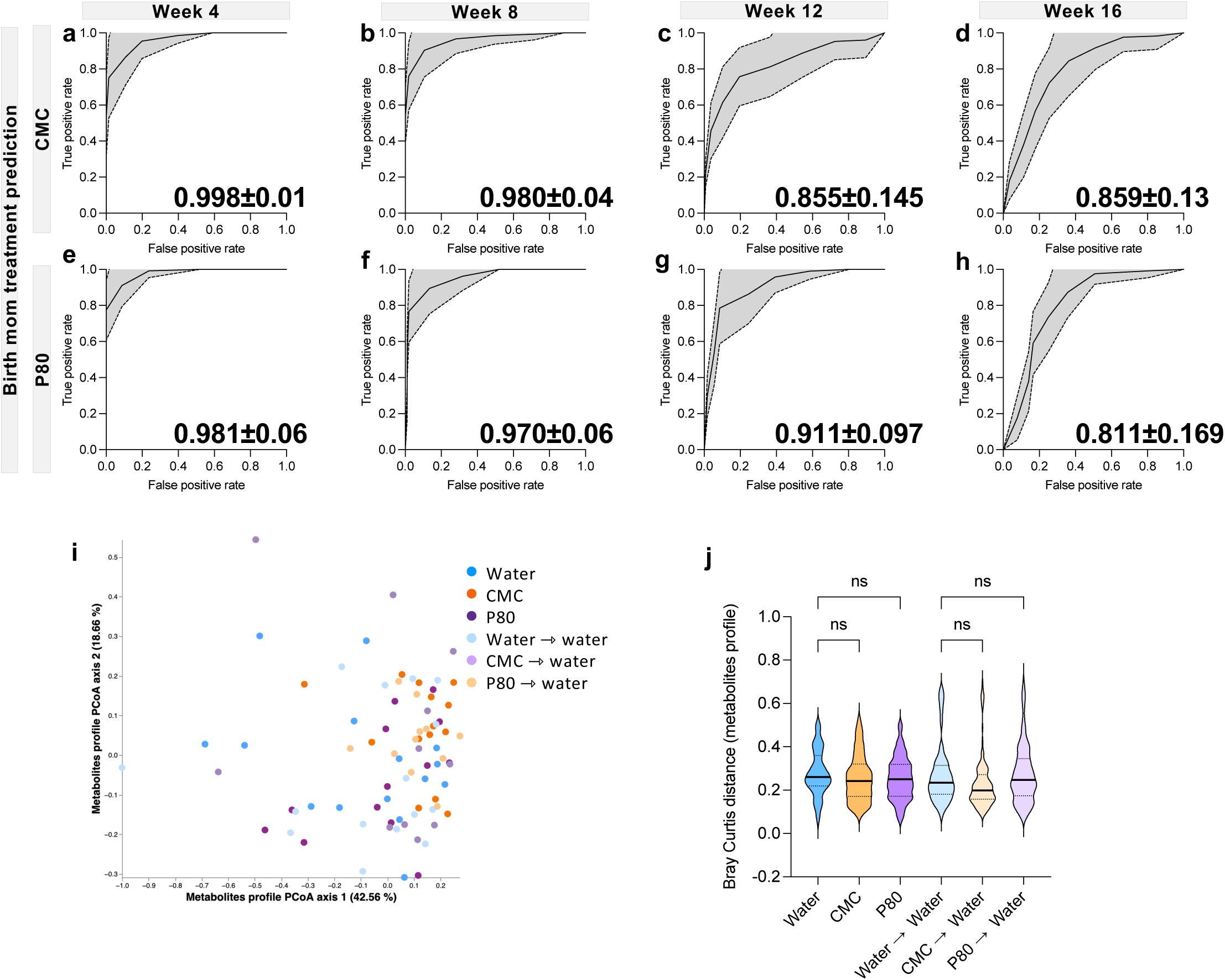
Maternal emulsifier intake results in early life-restricted microbiota alterations in offspring. Dams were subjected to 1% CMC or P80 in their drinking water for 10 weeks prior to breeding. At birth, offspring were either kept with their biological mother or cross-fostered to a water-treated dam. Following weaning of male offspring under emulsifier- free conditions, longitudinal microbiota composition and caecal metabolites were analyzed. (**a- h**) ROC curves-based prediction of maternal CMC treatment (outcome CMC or Water, **a-d**), or maternal P80 treatment (outcome P80 or Water, **e-h**), computed based on the rarefied abundance of offspring fecal microbiota members identified at the species level at week 4 (**a, e**), week 8 (**b, f**), week 12 (**c, g**) or week 16 (**d, h**). (**i, j**). Curves are mean ± s.d. of 50 predictive ROC iterations with independent training/validation cohort sampling, and mean area under the curve ± s.d. is indicated on the graph. Bray–Curtis distance was computed based on the relative abundance of 34 caecal metabolites quantified at week 16. PCoA plot of the Bray-Curtis distance (**i**) and Bray Curtis distance separating the various experimental groups from the water or water→water groups (**a-h**). N = 13-15 from ≥ 3 independent litters, significance was assessed by one-way ANOVA (**j**) and is indicated as follow: n.s., non-significant.

**Figure S3.**
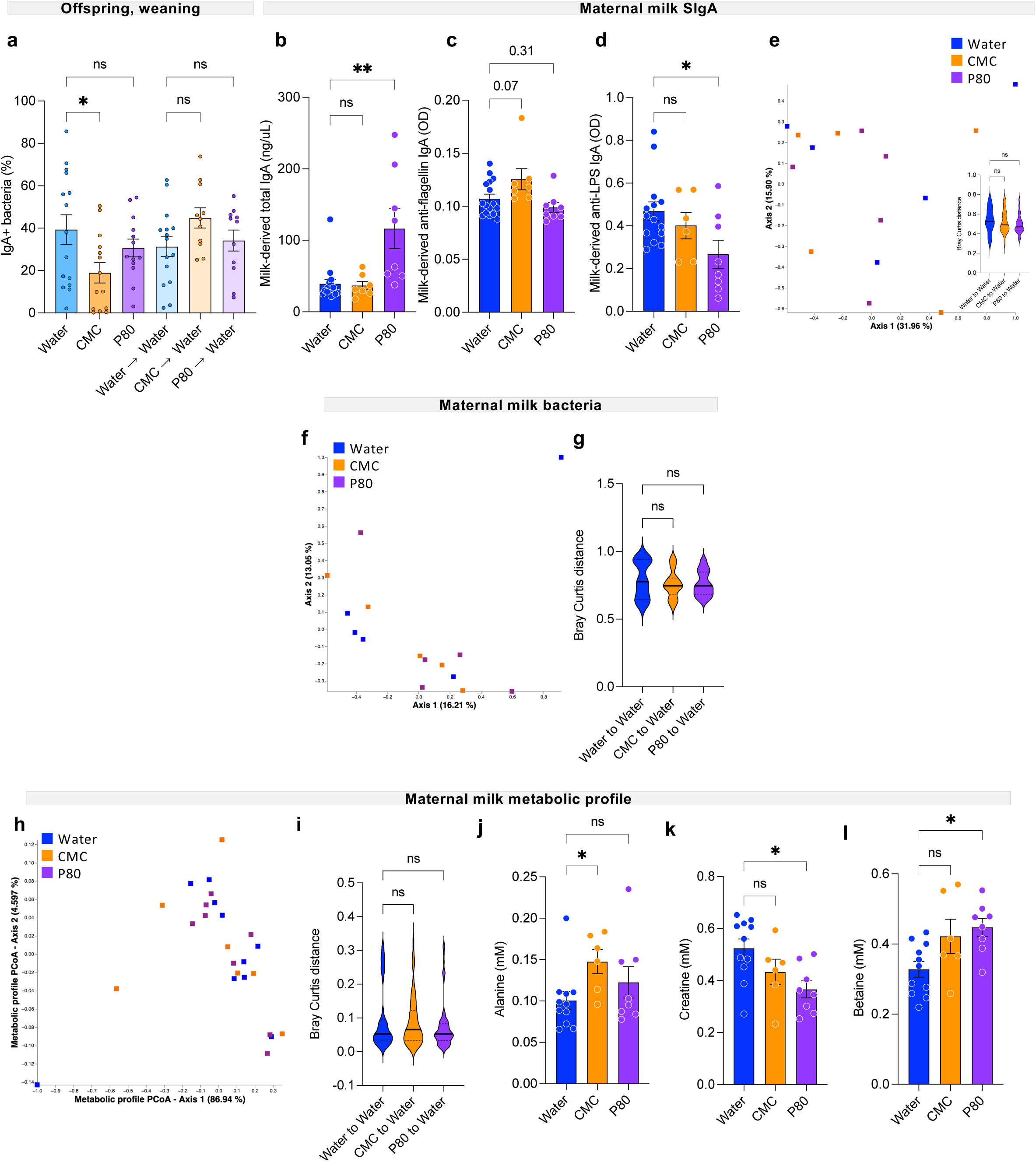
Maternal emulsifier intake is associated with modest modification in milk-derived components. Dams were subjected to 1% CMC or P80 in their drinking water for 10 weeks prior to breeding. At birth, offspring were either kept with their biological mother or cross-fostered to a water-treated dam. Dam’s milk was collected 14 days after birth. (**a**) Percentage of IgA-coated fecal bacteria in offspring fecal samples collected at weaning, assessed by flow cytometry. (**b**) Total milk IgA, (**c**) milk-derived anti-flagellin IgA and (**d**) anti- LPS IgA. (**e**) PCoA plot of the Bray Curtis distance of bacteria recognized by milk IgA, with the quantification of the Bray-Curtis distance separating the various experimental groups from the water-treated group. (**f-g**) Maternal milk resident bacteria were analyzed by 16S rRNA sequencing of milk pellets. (**f**) PCoA of the Bray Curtis distance of milk bacteria, and (**g**) Bray- Curtis distance separating the various experimental groups from the water-treated group. (**h-i**) Metabolites analysis of milk samples. (**h**) PCoA of the Bray Curtis distance computed based on the relative abundance of 17 milk metabolites, and (**i**) Bray-Curtis distance separating the various experimental groups from the water-treated group. (**j-k**) Absolute milk abundance of alanine (**j**), creatine (**k**) and betaine (**l**)). Data are the means ± s.e.m, N = 5-15. Significance was assessed by one-way ANOVA (**a-g, i-l**) and is indicated as follow: n.s., non-significant; * p≤ 0,05; **p ≤ 0,01.

**Figure S4.**
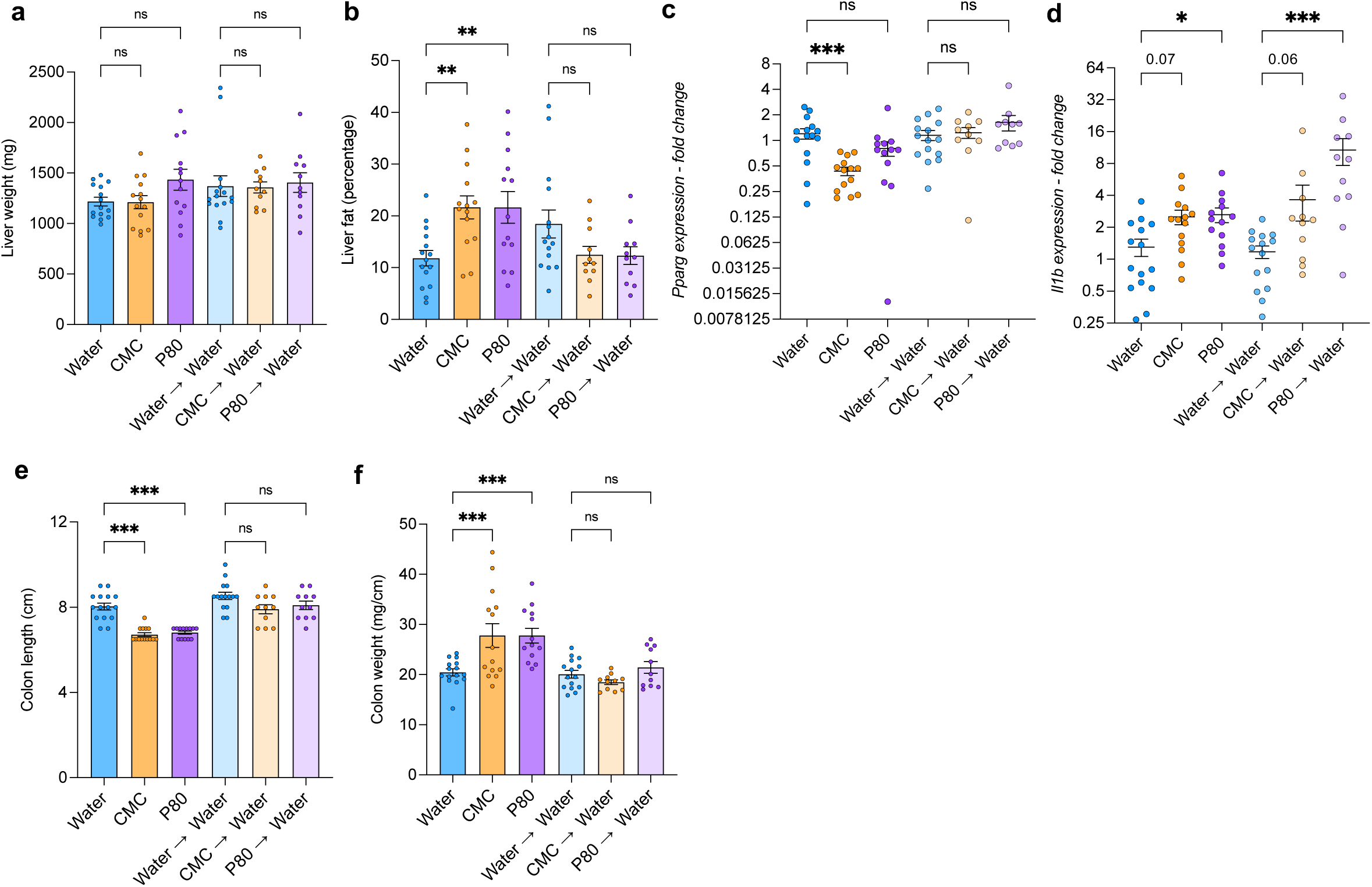
Maternal emulsifier intake increases offspring’ susceptibility to diet-induced obesity. Dams were subjected to 1% CMC or P80 in their drinking water for 10 weeks prior to breeding. At birth, offspring were either kept with their biological mother or cross- fostered to a water-treated dam. Following weaning of male offspring under emulsifier-free conditions, susceptibility to metabolic deregulations was assessed by subjecting mice to a HFD regimen for 13 weeks. (**a**) Liver weight, (**b**) liver fat measured as the average white area of 5 section of HE-stained liver. (**c-d**) Liver expression fold change of *Pparg* (**c**) or *Il1b* (**d**). (**e**) Colon length and (**f**) colon weight measured upon sacking. Data are the means ± s.e.m (N = 11- 15 from ≥ 3 independent litters). Significance was assessed by one-way ANOVA and is indicated as follow: n.s., nonsignificant; * p≤ 0,05; **p ≤ 0,01; ***p ≤ 0,001;.

**Figure S5.**
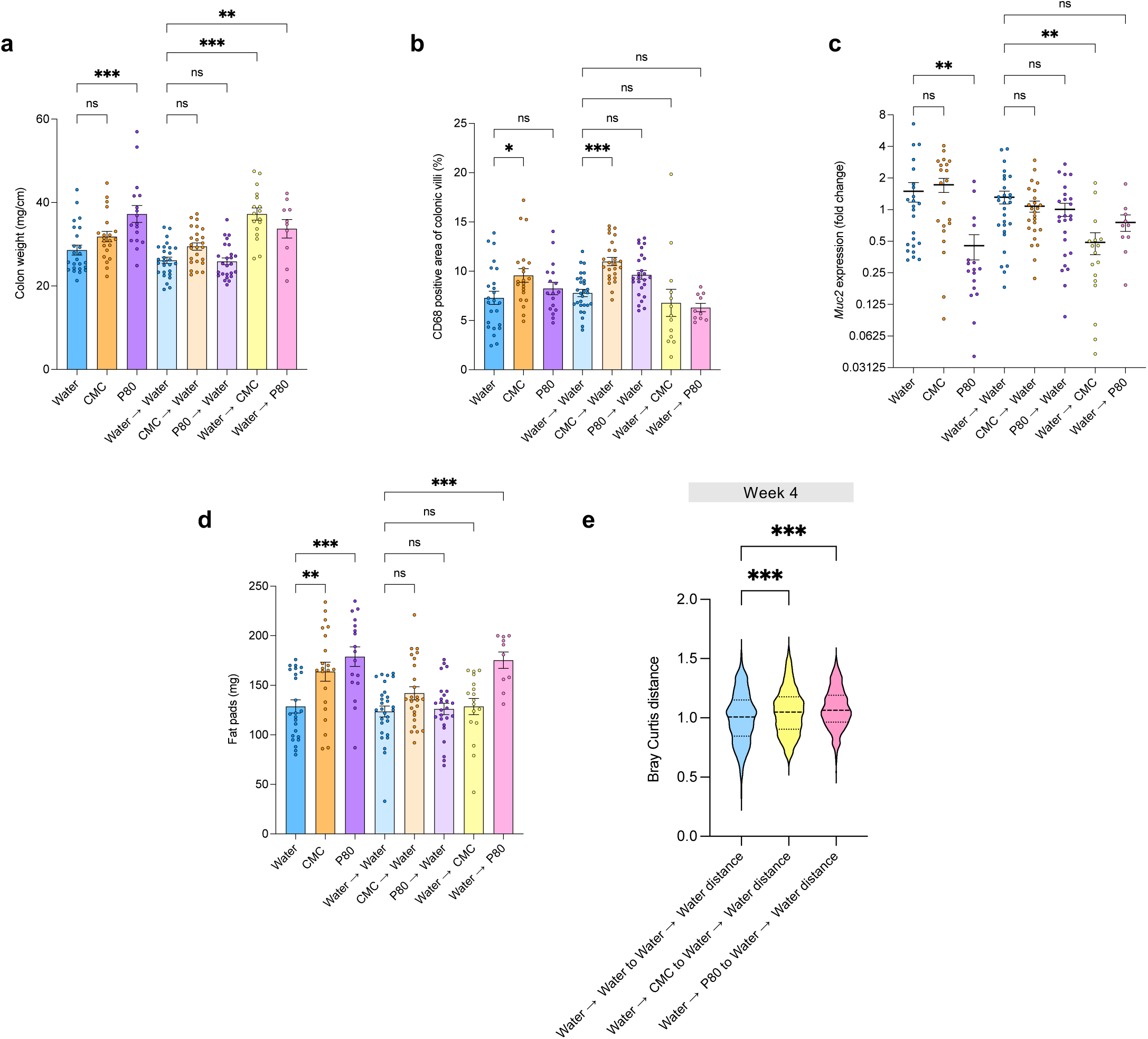
Maternal emulsifier intake increases offspring susceptibility to DSS- induced colitis. Dams were subjected to 1% CMC or P80 in their drinking water for 10 weeks prior to breeding. At birth, offspring were either kept with their biological mother or cross- fostered to a water-, CMC- or P80-treated dam. Male offspring were weaned under emulsifier- free condition, and susceptibility to intestinal inflammation was assessed by subjecting mice to a 1% DSS treatment at week 8 of age. (**a**) Colon weight upon sacking, (**b**) CD68^+^ cells infiltration quantified as the mean CD68 positive area per colonic crypt over 25 measurements, (**c**) colonic expression of *Muc2* and (**d**) epidydimal fat deposition measured upon sacking. (**e**) 16S microbiota composition represented as the Bray-Curtis distance distance separating the various experimental groups from the water→water group at week 4. Data are the means ± s.e.m (N = 10-28 from ≥ 3 independent litters). Significance was assessed by one-way ANOVA and is indicated as follow: n.s., nonsignificant, * p≤ 0,05; **p ≤ 0,01; ***p ≤ 0,001.

**Figure S6.**
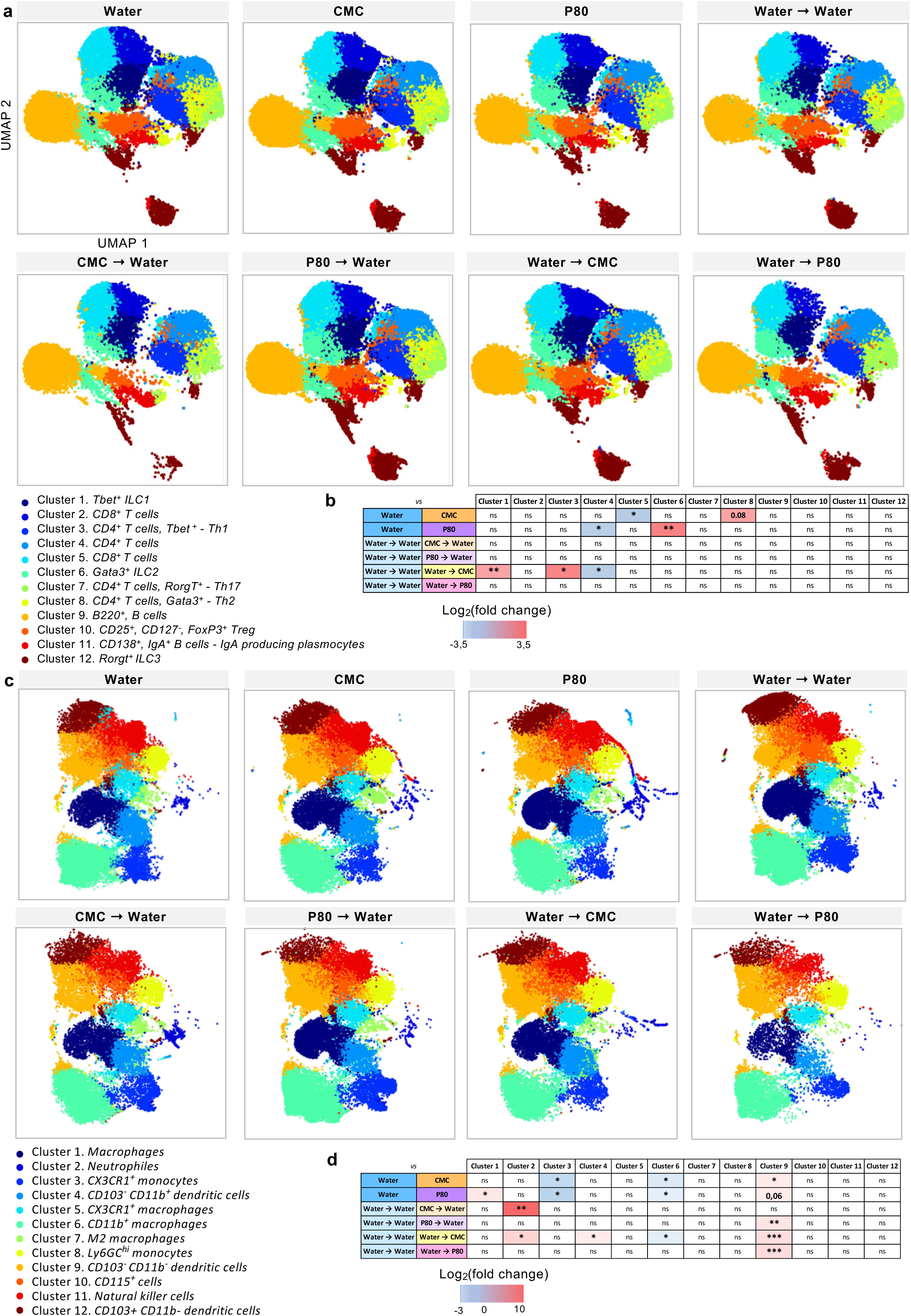
Maternal emulsifier intake results in alterations in immune populations following DSS-exposure. Dams were subjected to 1% CMC or P80 in their drinking water for 10 weeks prior to breeding. At birth, offspring were either kept with their biological mother or cross-fostered to a water-, CMC- or P80-treated dam. Male offspring were weaned under emulsifier-free condition, and susceptibility to intestinal inflammation was assessed by subjecting mice to a 1% DSS treatment at week 8 of age. Upon sacking, adaptive immune cells from the ileum lamina propria and innate immune cells from the colonic lamina propria were isolated and analyzed by flow cytometry. (**a**) UMAP computed on ileum-derived live CD45^+^,CD3^+^ and CD45+,B220^+^ cells, with unsupervised clustering computation. (**b**) Table of ileum immune cell clusters statistical comparison between the various experimental groups and the water or water→water offspring. (**c**) UMAP computed on colon-derived live CD45^+^ CD3^-^ B220^-^ with unsupervised clustering computation. (**c**) Table of colonic immune cell clusters statistical comparison between the various experimental groups and the water or water→water offspring. Significance was assessed by one-way ANOVA and is indicated as follow n.s. indicates nonsignificant; * p≤ 0,05; **p ≤ 0,01; ***p ≤ 0,001.

**Figure S7.**
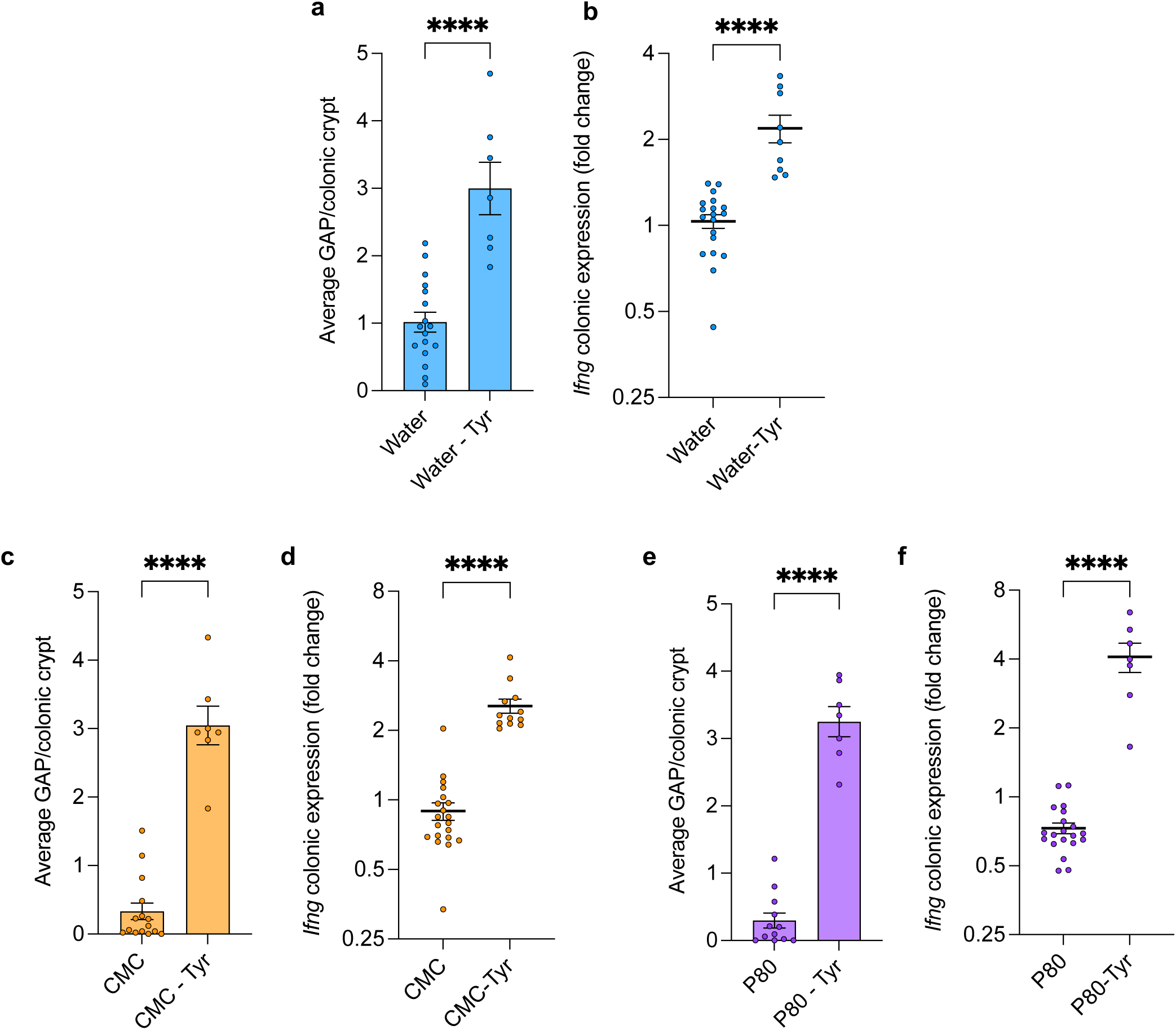
Colonic goblet cell-associated passages opening levels associates with *Ifng* expression. Dams were subjected to 1% CMC or P80 in their drinking water for 10 weeks prior to breeding, and obtained pups were either left untreated or daily injected, from day 10 to day 21, with tyrphostin in order to maintain colonic goblet-cell associated antigens passages (GAPs) opening. At weaning, GAPs opening levels and *Ifng* colonic expression were assessed. (**a-f**) Quantification of GAPs and *Ifng* colonic expression fold change in tyrphostin injected or untreated water offspring (**a, b**), in tyrphostin injected or untreated CMC offspring (**c, d**) or in tyrphostin injected or untreated P80 offspring (**e, f**). Data are the means ± s.e.m (N = 10-28 from ≥ 3 independent litters). Significance was assessed by one-way ANOVA and is indicated as **** p ≤ 0,0001.

**Figure S8.**
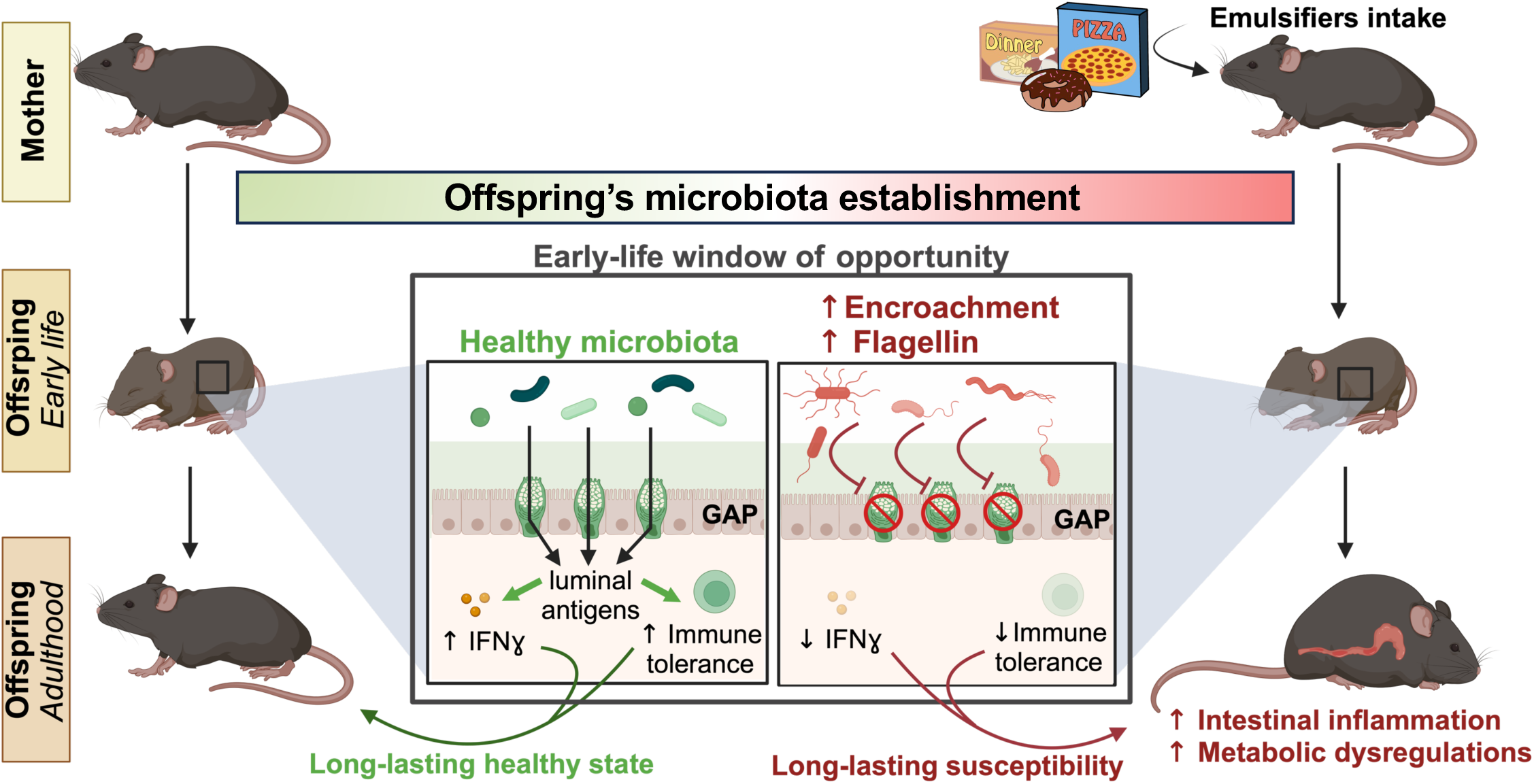
Schematic outline of the main findings.

## Notes

Conflict of interest: **none.**

### Competing Interest Statement

The authors have declared no competing interest.

## References

1. Al Nabhani, Z., Dulauroy, S., Marques, R., Cousu, C., Al Bounny, S., Déjardin, F., Sparwasser, T., Bérard, M., Cerf-Bensussan, N., and Eberl, G. (2019). A Weaning Reaction to Microbiota Is Required for Resistance to Immunopathologies in the Adult. Immunity 50, 1276–1288.e5. 10.1016/j.immuni.2019.02.014.

2. Cox, L.M., Yamanishi, S., Sohn, J., Alekseyenko, A.V., Leung, J.M., Cho, I., Kim, S.G., Li, H., Gao, Z., Mahana, D., et al. (2014). Altering the Intestinal Microbiota during a Critical Developmental Window Has Lasting Metabolic Consequences. Cell 158, 705–721. 10.1016/j.cell.2014.05.052.

3. Hornef, M.W., and Torow, N. (2020). ‘Layered immunity’ and the ‘neonatal window of opportunity’ – timed succession of non-redundant phases to establish mucosal host– microbial homeostasis after birth. Immunology 159, 15–25. 10.1111/imm.13149.

4. El Aidy, S., Hooiveld, G., Tremaroli, V., Bäckhed, F., and Kleerebezem, M. (2013). The gut microbiota and mucosal homeostasis: Colonized at birth or at adulthood, does it matter? Gut Microbes 4, 118–124. 10.4161/gmic.23362.

5. Gensollen, T., Iyer, S.S., Kasper, D.L., and Blumberg, R.S. (2016). How colonization by microbiota in early life shapes the immune system. Science 352, 539–544. 10.1126/science.aad9378.

6. Olszak, T., An, D., Zeissig, S., Vera, M.P., Richter, J., Franke, A., Glickman, J.N., Siebert, R., Baron, R.M., Kasper, D.L., et al. (2012). Microbial Exposure During Early Life Has Persistent Effects on Natural Killer T Cell Function. Science 336, 489–493. 10.1126/science.1219328.

7. Smith, P.M., Howitt, M.R., Panikov, N., Michaud, M., Gallini, C.A., Bohlooly-Y, M., Glickman, J.N., and Garrett, W.S. (2013). The microbial metabolites, short chain fatty acids, regulate colonic Treg cell homeostasis. Science 341, 10.1126/science.1241165. https://doi.org/10.1126/science.1241165.

8. Constantinides, M.G., Link, V.M., Tamoutounour, S., Wong, A.C., Perez-Chaparro, P.J., Han, S.-J., Chen, Y.E., Li, K., Farhat, S., Weckel, A., et al. (2019). MAIT cells are imprinted by the microbiota in early life and promote tissue repair. Science 366, eaax6624. 10.1126/science.aax6624.

9. Al Nabhani, Z., Dulauroy, S., Lécuyer, E., Polomack, B., Campagne, P., Berard, M., and Eberl, G. (2019). Excess calorie intake early in life increases susceptibility to colitis in adulthood. Nat Metab 1, 1101–1109. 10.1038/s42255-019-0129-5.

10. Scheer, S., Medina, T.S., Murison, A., Taves, M.D., Antignano, F., Chenery, A., Soma, K.K., Perona-Wright, G., Lupien, M., Arrowsmith, C.H., et al. (2017). Early-life antibiotic treatment enhances the pathogenicity of CD4 ^+^ T cells during intestinal inflammation. J Leukoc Biol 101, 893–900. 10.1189/jlb.3MA0716-334RR.

11. Adami, A.J., Bracken, S.J., Guernsey, L.A., Rafti, E., Maas, K.R., Graf, J., Matson, A.P., Thrall, R.S., and Schramm, C.M. (2018). Early-life antibiotics attenuate regulatory T cell generation and increase the severity of murine house dust mite-induced asthma. Pediatr Res 84, 426–434. 10.1038/s41390-018-0031-y.

12. Russell, S.L., Gold, M.J., Hartmann, M., Willing, B.P., Thorson, L., Wlodarska, M., Gill, N., Blanchet, M., Mohn, W.W., McNagny, K.M., et al. (2012). Early life antibiotic-driven changes in microbiota enhance susceptibility to allergic asthma. EMBO Rep 13, 440–447. 10.1038/embor.2012.32.

13. Cho, I., Yamanishi, S., Cox, L., Methé, B.A., Zavadil, J., Li, K., Gao, Z., Mahana, D., Raju, K., Teitler, I., et al. (2012). Antibiotics in early life alter the murine colonic microbiome and adiposity. Nature 488, 621–626. 10.1038/nature11400.

14. Zou, J., Ngo, V.L., Wang, Y., Wang, Y., and Gewirtz, A.T. (2023). Maternal fiber deprivation alters microbiota in offspring, resulting in low-grade inflammation and predisposition to obesity. Cell Host & Microbe 31, 45–57.e7. 10.1016/j.chom.2022.10.014.

15. Shelton, C.D., Sing, E., Mo, J., Shealy, N.G., Yoo, W., Thomas, J., Fitz, G.N., Castro, P.R., Hickman, T.T., Torres, T.P., et al. (2023). An early-life microbiota metabolite protects against obesity by regulating intestinal lipid metabolism. Cell Host & Microbe 31, 1604–1619.e10. 10.1016/j.chom.2023.09.002.

16. Knoop, K.A., Gustafsson, J.K., McDonald, K.G., Kulkarni, D.H., Coughlin, P.E., McCrate, S., Kim, D., Hsieh, C.-S., Hogan, S.P., Elson, C.O., et al. (2017). Microbial antigen encounter during a preweaning interval is critical for tolerance to gut bacteria. Sci. Immunol. 2, eaao1314. 10.1126/sciimmunol.aao1314.

17. Knoop, K.A., Gustafsson, J.K., McDonald, K.G., Kulkarni, D.H., Kassel, R., and Newberry, R.D. (2017). Antibiotics promote the sampling of luminal antigens and bacteria via colonic goblet cell associated antigen passages. Gut Microbes 8, 400–411. 10.1080/19490976.2017.1299846.

18. Knoop, K.A., McDonald, K.G., McCrate, S., McDole, J.R., and Newberry, R.D. (2015). Microbial sensing by goblet cells controls immune surveillance of luminal antigens in the colon. Mucosal Immunol 8, 198–210. 10.1038/mi.2014.58.

19. Kulkarni, D.H., Gustafsson, J.K., Knoop, K.A., McDonald, K.G., Bidani, S.S., Davis, J.E., Floyd, A.N., Hogan, S.P., Hsieh, C.-S., and Newberry, R.D. (2020). Goblet cell associated antigen passages support the induction and maintenance of oral tolerance. Mucosal Immunology 13, 271–282. 10.1038/s41385-019-0240-7.

20. Schulfer, A.F., Battaglia, T., Alvarez, Y., Bijnens, L., Ruiz, V.E., Ho, M., Robinson, S., Ward, T., Cox, L.M., Rogers, A.B., et al. (2018). Intergenerational transfer of antibiotic-perturbed microbiota enhances colitis in susceptible mice. Nat Microbiol 3, 234–242. 10.1038/s41564-017-0075-5.

21. Valles-Colomer, M., Blanco-Míguez, A., Manghi, P., Asnicar, F., Dubois, L., Golzato, D., Armanini, F., Cumbo, F., Huang, K.D., Manara, S., et al. (2023). The person-to-person transmission landscape of the gut and oral microbiomes. Nature 614, 125–135. 10.1038/s41586-022-05620-1.

22. Valles-Colomer, M., Bacigalupe, R., Vieira-Silva, S., Suzuki, S., Darzi, Y., Tito, R.Y., Yamada, T., Segata, N., Raes, J., and Falony, G. (2021). Variation and transmission of the human gut microbiota across multiple familial generations. Nat Microbiol 7, 87–96. 10.1038/s41564-021-01021-8.

23. Gopalakrishna, K.P., and Hand, T.W. (2020). Influence of Maternal Milk on the Neonatal Intestinal Microbiome. Nutrients 12, 823. 10.3390/nu12030823.

24. Rogier, E.W., Frantz, A.L., Bruno, M.E.C., Wedlund, L., Cohen, D.A., Stromberg, A.J., and Kaetzel, C.S. (2014). Secretory antibodies in breast milk promote long-term intestinal homeostasis by regulating the gut microbiota and host gene expression. Proc. Natl. Acad. Sci. U.S.A. 111, 3074–3079. 10.1073/pnas.1315792111.

25. Kalbermatter, C., Fernandez Trigo, N., Christensen, S., and Ganal-Vonarburg, S.C. (2021). Maternal Microbiota, Early Life Colonization and Breast Milk Drive Immune Development in the Newborn. Front. Immunol. 12, 683022. 10.3389/fimmu.2021.683022.

26. Chassaing, B., Koren, O., Goodrich, J.K., Poole, A.C., Srinivasan, S., Ley, R.E., and Gewirtz, A.T. (2015). Dietary emulsifiers impact the mouse gut microbiota promoting colitis and metabolic syndrome. Nature 519, 92–96. 10.1038/nature14232.

27. Chassaing, B., Van de Wiele, T., De Bodt, J., Marzorati, M., and Gewirtz, A.T. (2017). Dietary emulsifiers directly alter human microbiota composition and gene expression ex vivo potentiating intestinal inflammation. Gut 66, 1414–1427. 10.1136/gutjnl-2016-313099.

28. Chassaing, B., Compher, C., Bonhomme, B., Liu, Q., Tian, Y., Walters, W., Nessel, L., Delaroque, C., Hao, F., Gershuni, V., et al. (2021). Randomized controlled-feeding study of dietary emulsifier carboxymethylcellulose reveals detrimental impacts on the gut microbiota and metabolome. Gastroenterology, S0016-5085(21)03728-8. 10.1053/j.gastro.2021.11.006.

29. Delaroque, C., and Chassaing, B. (2024). Dietary emulsifier consumption accelerates type 1 diabetes development in NOD mice. NPJ Biofilms Microbiomes 10, 1. 10.1038/s41522-023-00475-4.

30. Salame, C., Javaux, G., Sellem, L., Viennois, E., de Edelenyi, F.S., Agaësse, C., De Sa, A., Huybrechts, I., Pierre, F., Coumoul, X., et al. (2024). Food additive emulsifiers and the risk of type 2 diabetes: analysis of data from the NutriNet-Santé prospective cohort study. Lancet Diabetes Endocrinol 12, 339–349. 10.1016/S2213-8587(24)00086-X.

31. Sellem, L., Srour, B., Javaux, G., Chazelas, E., Chassaing, B., Viennois, E., Debras, C., Salamé, C., Druesne-Pecollo, N., Esseddik, Y., et al. (2023). Food additive emulsifiers and risk of cardiovascular disease in the NutriNet-Santé cohort: prospective cohort study. BMJ, e076058. 10.1136/bmj-2023-076058.

32. Sellem, L., Srour, B., Javaux, G., Chazelas, E., Chassaing, B., Viennois, E., Debras, C., Druesne-Pecollo, N., Esseddik, Y., Szabo de Edelenyi, F., et al. (2024). Food additive emulsifiers and cancer risk: Results from the French prospective NutriNet-Santé cohort. PLoS Med 21, e1004338. 10.1371/journal.pmed.1004338.

33. Holst, A.Q., Jois, H., Laursen, M.F., Sommer, M.O.A., Licht, T.R., and Bahl, M.I. (2022). Human milk oligosaccharides induce acute yet reversible compositional changes in the gut microbiota of conventional mice linked to a reduction of butyrate levels. Microlife 3, uqac006. 10.1093/femsml/uqac006.

34. Boudry, G., Charton, E., Le Huerou-Luron, I., Ferret-Bernard, S., Le Gall, S., Even, S., and Blat, S. (2021). The Relationship Between Breast Milk Components and the Infant Gut Microbiota. Front Nutr 8, 629740. 10.3389/fnut.2021.629740.

35. Vijay-Kumar, M., Bovilla, V.R., Yeoh, B.S., Golonka, R.M., Saha, P., Joe, B., and Gewirtz, A.T. (2023). Bacterial flagellin is a dominant, stable innate immune activator in the gastrointestinal contents of mice and rats. Gut Microbes 15, 2185031. 10.1080/19490976.2023.2185031.

36. Chassaing, B., Koren, O., Carvalho, F.A., Ley, R.E., and Gewirtz, A.T. (2014). AIEC pathobiont instigates chronic colitis in susceptible hosts by altering microbiota composition. Gut 63, 1069–1080. 10.1136/gutjnl-2013-304909.

37. Naimi, S., Viennois, E., Gewirtz, A.T., and Chassaing, B. (2021). Direct impact of commonly used dietary emulsifiers on human gut microbiota. Microbiome 9, 66. 10.1186/s40168-020-00996-6.

38. Chassaing, B., Raja, S.M., Lewis, J.D., Srinivasan, S., and Gewirtz, A.T. (2017). Colonic Microbiota Encroachment Correlates With Dysglycemia in Humans. Cellular and Molecular Gastroenterology and Hepatology 4, 205–221. 10.1016/j.jcmgh.2017.04.001.

39. Johansson, M.E.V., Gustafsson, J.K., Holmén-Larsson, J., Jabbar, K.S., Xia, L., Xu, H., Ghishan, F.K., Carvalho, F.A., Gewirtz, A.T., Sjövall, H., et al. (2014). Bacteria penetrate the normally impenetrable inner colon mucus layer in both murine colitis models and patients with ulcerative colitis. Gut 63, 281–291. 10.1136/gutjnl-2012-303207.

40. Sevrin, G., Massier, S., Chassaing, B., Agus, A., Delmas, J., Denizot, J., Billard, E., and Barnich, N. (2020). Adaptation of adherent-invasive E. coli to gut environment: Impact on flagellum expression and bacterial colonization ability. Gut Microbes 11, 364–380. 10.1080/19490976.2017.1421886.

41. Gulhane, M., Murray, L., Lourie, R., Tong, H., Sheng, Y.H., Wang, R., Kang, A., Schreiber, V., Wong, K.Y., Magor, G., et al. (2016). High Fat Diets Induce Colonic Epithelial Cell Stress and Inflammation that is Reversed by IL-22. Sci Rep 6, 28990. 10.1038/srep28990.

42. Torow, N., and Hornef, M.W. (2017). The Neonatal Window of Opportunity: Setting the Stage for Life-Long Host-Microbial Interaction and Immune Homeostasis. J.I. 198, 557–563. 10.4049/jimmunol.1601253.

43. Sefik, E., Geva-Zatorsky, N., Oh, S., Cahenzli, J., Köller, Y., Wyss, M., Geuking, M.B., McCoy, K.D., Mathis, D., and Benoist, C. Individual intestinal symbionts induce a distinct population of RORg + regulatory T cells. 6.

44. Viennois, E., Merlin, D., Gewirtz, A.T., and Chassaing, B. (2017). Dietary Emulsifier– Induced Low-Grade Inflammation Promotes Colon Carcinogenesis. Cancer Res 77, 27–40. 10.1158/0008-5472.CAN-16-1359.

45. Fulde, M., Sommer, F., Chassaing, B., van Vorst, K., Dupont, A., Hensel, M., Basic, M., Klopfleisch, R., Rosenstiel, P., Bleich, A., et al. (2018). Neonatal selection by Toll-like receptor 5 influences long-term gut microbiota composition. Nature 560, 489–493. 10.1038/s41586-018-0395-5.

46. Ganal-Vonarburg, S.C., Fuhrer, T., and Gomez de Agüero, M. (2017). Maternal microbiota and antibodies as advocates of neonatal health. Gut Microbes 8, 479–485. 10.1080/19490976.2017.1299847.

47. Liang, W., Menke, A.L., Driessen, A., Koek, G.H., Lindeman, J.H., Stoop, R., Havekes, L.M., Kleemann, R., and van den Hoek, A.M. (2014). Establishment of a General NAFLD Scoring System for Rodent Models and Comparison to Human Liver Pathology. PLoS One 9, e115922. 10.1371/journal.pone.0115922.

48. Chassaing, B., Srinivasan, G., Delgado, M.A., Young, A.N., Gewirtz, A.T., and Vijay- Kumar, M. (2012). Fecal Lipocalin 2, a Sensitive and Broadly Dynamic Non-Invasive Biomarker for Intestinal Inflammation. PLoS ONE 7, e44328. 10.1371/journal.pone.0044328.

49. Martin, M. (2011). Cutadapt removes adapter sequences from high-throughput sequencing reads. EMBnet.journal 17, 10–12. 10.14806/ej.17.1.200.

50. Langmead, B., and Salzberg, S.L. (2012). Fast gapped-read alignment with Bowtie 2. Nat Methods 9, 357–359. 10.1038/nmeth.1923.

51. Trapnell, C., Roberts, A., Goff, L., Pertea, G., Kim, D., Kelley, D.R., Pimentel, H., Salzberg, S.L., Rinn, J.L., and Pachter, L. (2012). Differential gene and transcript expression analysis of RNA-seq experiments with TopHat and Cufflinks. Nat Protoc 7, 562–578. 10.1038/nprot.2012.016.

52. Robinson, M.D., McCarthy, D.J., and Smyth, G.K. (2010). edgeR : a Bioconductor package for differential expression analysis of digital gene expression data. Bioinformatics 26, 139–140. 10.1093/bioinformatics/btp616.

53. Mi, H., Muruganujan, A., Casagrande, J.T., and Thomas, P.D. (2013). Large-scale gene function analysis with the PANTHER classification system. Nat Protoc 8, 1551–1566. 10.1038/nprot.2013.092.

54. Smith, L., Klément, W., Dopavogui, L., de Bock, F., Lasserre, F., Barretto, S., Lukowicz, C., Fougerat, A., Polizzi, A., Schaal, B., et al. (2020). Perinatal exposure to a dietary pesticide cocktail does not increase susceptibility to high-fat diet-induced metabolic perturbations at adulthood but modifies urinary and fecal metabolic fingerprints in C57Bl6/J mice. Environment International 144, 106010. 10.1016/j.envint.2020.106010.

55. Matamoros, C., Hao, F., Tian, Y., Patterson, A.D., and Harvatine, K.J. (2022). Interaction of sodium acetate supplementation and dietary fiber level on feeding behavior, digestibility, milk synthesis, and plasma metabolites. J Dairy Sci 105, 8824–8838. 10.3168/jds.2022-21911.

56. Tran, H.Q., Ley, R.E., Gewirtz, A.T., and Chassaing, B. (2019). Flagellin-elicited adaptive immunity suppresses flagellated microbiota and vaccinates against chronic inflammatory diseases. Nat Commun 10, 5650. 10.1038/s41467-019-13538-y.

57. Caporaso, J.G., Lauber, C.L., Walters, W.A., Berg-Lyons, D., Huntley, J., Fierer, N., Owens, S.M., Betley, J., Fraser, L., Bauer, M., et al. (2012). Ultra-high-throughput microbial community analysis on the Illumina HiSeq and MiSeq platforms. ISME J 6, 1621–1624. 10.1038/ismej.2012.8.

58. Bolyen, E., Rideout, J.R., Dillon, M.R., Bokulich, N.A., Abnet, C.C., Al-Ghalith, G.A., Alexander, H., Alm, E.J., Arumugam, M., Asnicar, F., et al. (2019). Reproducible, interactive, scalable and extensible microbiome data science using QIIME 2. Nat Biotechnol 37, 852–857. 10.1038/s41587-019-0209-9.

59. Callahan, B.J., McMurdie, P.J., Rosen, M.J., Han, A.W., Johnson, A.J.A., and Holmes, S.P. (2016). DADA2: High-resolution sample inference from Illumina amplicon data. Nat Methods 13, 581–583. 10.1038/nmeth.3869.

60. Mallick, H., Rahnavard, A., McIver, L.J., Ma, S., Zhang, Y., Nguyen, L.H., Tickle, T.L., Weingart, G., Ren, B., Schwager, E.H., et al. (2021). Multivariable association discovery in population-scale meta-omics studies. PLoS Comput Biol 17, e1009442. 10.1371/journal.pcbi.1009442.

61. Mombaerts, P., Iacomini, J., Johnson, R.S., Herrup, K., Tonegawa, S., and Papaioannou, V.E. (1992). RAG-1-deficient mice have no mature B and T lymphocytes. Cell 68, 869– 877. 10.1016/0092-8674(92)90030-G.

62. Johansson, M.E.V., and Hansson, G.C. (2012). Preservation of mucus in histological sections, immunostaining of mucins in fixed tissue, and localization of bacteria with FISH. Methods Mol Biol., 229–235. 10.1007/978-1-61779-513-8_13.

